# Breast Cancer-derived PDGFB Remodels the CNS Vascular Niche to Promote Brain, Spinal and Leptomeningeal Metastasis

**DOI:** 10.64898/2026.10.15.759418

**Authors:** Nathaniel S. Grabinski, Jesse J. Reardon, Sajita Shah, Jonathan J. Adorno, Paul Lee, Rebecca L. Packard, Yalini Ramamoorthy, Katie A. Thies, Prathik A. Chakravarthy, Shashwat S. Agarwal, Jonathan M. Spehar, Tasneem A. Arsiwala, James W. Carpenter, Alexis A. Mossing, Johnathon G. Schiebel, Sarah A. Steck, Alexander J. Didier, Dillon S. Richardson, Luke Kollin, Monica Venere, Jonathan P. Godbout, Paul R. Lockman, Sasha Beyer, Kara N. Corps, Jessica O. Winter, Jonathan W. Song, Steven T. Sizemore, Gina M. Sizemore

**Affiliations:** Department of Radiation Oncology, The Ohio State University Wexner Medical Center, Columbus, Ohio 43210, USA; The Comprehensive Cancer Center, The Ohio State University, Columbus, Ohio 43210, USA; Neuroscience Graduate Program, The Ohio State University, Columbus, OH 43210, USA; Department of Biomedical Engineering, The Ohio State University, Columbus, Ohio 43210, USA; William G. Lowrie Department of Chemical and Biomolecular Engineering, The Ohio State University, Columbus, Ohio 43210, USA; Department of Mechanical and Aerospace Engineering, The Ohio State University, Columbus, Ohio 43210, USA; Rockefeller Neuroscience Institute, West Virginia University, Morgantown, WV 26506, USA; Department of Neuroscience, The Ohio State University, Columbus, Ohio 43210, USA; Center for Brain Injury Recovery & Discovery, The Ohio State University, Columbus, Ohio 43210, USA; Department of Basic Pharmaceutical Sciences, West Virginia University School of Pharmacy, Morgantown, WV 26506, USA; Department of Veterinary Biosciences, The Ohio State University, Columbus, Ohio 43210, USA

**Author notes:** Correspondence: Gina M. Sizemore, Ph.D. Associate Professor, Department of Radiation Oncology, College of Medicine, Cancer Biology Program Co-Leader, The Ohio State University Comprehensive Cancer Center The Ohio State University, 646C Tzagournis Medical Research Facility, 420 W. 12th Avenue, Columbus, OH 43210. co-first authorship (Nathaniel S. Grabinski, Jesse J. Reardon, Sajita Shah).

**Keywords:** Breast cancer, CNS metastasis, leptomeningeal disease, PDGFB, angiogenesis

## Abstract

**Background:** Central nervous system (CNS) metastases are a major driver of breast cancer (BC) mortality. Recently, our group demonstrated that platelet-derived growth factor-B (PDGFB) and its receptor, platelet-derived growth factor receptor-beta (PDGFRβ), promote BC metastasis to the brain; however, the cellular/molecular mechanisms driving this process remain unclear. Here, we investigated tumor-derived PDGFB effects on the CNS vascular niche in regulating metastatic spread.

**Methods:** Triple-negative BC cells were engineered to overexpress PDGFB and used to evaluate endothelial sprouting, blood-brain barrier (BBB) function, vascular permeability and experimental CNS metastasis. A novel retro-orbital (RO) injection model was used to study leptomeningeal/spinal experimental metastasis.

**Results:** PDGFB enhanced human brain microvascular endothelial cell (HBMEC) sprouting, induced cell migration in BBB spheroids, increased permeability in 3D models, and disrupted tight junction proteins in HBMECs. Notably, alterations in vascular permeability were observed in both PDGFRβ-positive and PDGFRβ-negative endothelial systems, suggesting PDGFB can remodel the CNS vasculature through multiple mechanisms. *In vivo*, PDGFB increased CNS metastatic burden and altered tumor vasculature in xenograft and allograft models of BC. Furthermore, PDGFB promoted robust colonization of both spinal and leptomeningeal compartments, identifying a previously underappreciated role for this signaling pathway in a clinically devastating form of CNS metastasis. Analysis of patient transcriptomic datasets revealed enrichment of angiogenesis-related pathways in PDGFB-high tumors from patients who developed CNS metastases.

**Conclusions:** Collectively, these findings identify tumor-derived PDGFB as a key regulator of CNS vascular remodeling and metastatic tropism, while introducing a new experimental framework to study spinal metastases and leptomeningeal disease in BC.

**Key Points:**

- Tumor-derived PDGFB remodels the CNS vasculature and disrupts barrier integrity
- PDGFB promotes brain, spinal, and leptomeningeal metastasis
- Retro-orbital injections provide a new model of leptomeningeal and spinal metastasis

**Importance of Study:** Central nervous system (CNS) metastases are among the most lethal complications of breast cancer. Vascular cooption is recognized as a critical step in the metastatic cascade, but the tumor-derived signals which mediate this process are not fully understood. This study identifies PDGFB as a key signaling molecule which modulates the CNS endothelium, a previously underappreciated function. Beyond advancing our understanding of PDGFB-driven brain metastasis, this work also expands the role of PDGFB in promoting spinal and leptomeningeal metastasis. We establish a mechanistic framework connecting tumor-derived PDGFB to vascular dysfunction and CNS tropism, highlighting a potential opportunity to improve patient prognosis. Additionally, we employ a novel experimental model of leptomeningeal disease through retro-orbital (RO) injection of breast cancer cells. Together, these findings provide new insight into CNS metastasis while establishing experimental tools that may accelerate advancements in the understanding and treatment of brain, spinal, and leptomeningeal metastasis.

## Introduction

In 2026, breast cancer (BC) was the most commonly diagnosed cancer in women in the world with 321,910 new cases this year^1^, and despite significant advancements in the field, metastasis remains the major cause of BC-related mortality^2^. Breast cancer-associated central nervous system (BC-CNS) metastasis, consisting of parenchymal metastases and leptomeningeal disease (LMD), represent particularly devastating complications of advanced BC^3^. Depending on the molecular subtype, up to 30-50% of patients with HER2-positive or triple-negative (TN) disease will ultimately develop CNS involvement^4–7^. Breast cancer-associated leptomeningeal disease (BC-LMD) affects 5-20% of metastatic BC patients^3^, and those with TN disease account for the highest share of cases relative to its prevalence in the overall BC population^8,9^. Once diagnosed with CNS metastases, patient outcomes decrease significantly. Median survival for patients with BC-associated brain metastasis (BCBM) is ∼10-11 months^10^, whereas patients diagnosed with BC-LMD frequently only live 3-6 months, with TNBC patients faring the worst^9,11^. In comparison, patients developing lung, liver and bone metastases have median survival rates approaching 30 to 40 months^12,13^. This unfavorable outcome of BC-CNS metastasis is in part attributed to the inability of treatment options to traverse the blood-brain barrier (BBB)^14^.

Improving outcomes for patients with BC-CNS metastases requires a better understanding of the intricate molecular mechanisms that drive metastatic colonization and progression in the CNS. A continued knowledge gap is the functional interplay between BC cells and the highly complex metastatic CNS tumor microenvironment (TME) that is extremely different than other sites of BC metastasis^15^. This site harbors unique cellular components found only in the CNS including astrocytes, microglia, oligodendrocytes, and neurons. The CNS TME also contains a hyaluronic acid rich extracellular matrix (ECM), which is specific to the brain^16^. Additionally, the brain is a unique site for metastasis due to the presence of the BBB and the blood-CSF barrier (BCSFB), which serve as CNS gatekeepers. The BBB and BCSFB shield the brain from potentially harmful pathogens and toxins^17,18^. This protection is achieved through tight junctions that seal the endothelial cells of brain capillaries in the BBB or the leptomeningeal cells and specialized epithelial cells in the BCSFB^19–22^. Furthermore, the cerebrospinal fluid (CSF) filled leptomeningeal space dynamically interacts with the entire brain, where it functions in homeostasis and waste clearance through the glymphatic system^23^. Thus, the presence of LMD can rapidly impair the whole CNS, highlighting the importance of identifying novel mechanisms by which BC cells adapt and survive in this unique space.

Successful metastatic spread requires circulating tumor cells (CTCs) to cross the BBB and/or BCSFB via trans-endothelial migration^24^ through various mechanisms including adhesion, degradation of the ECM, and modification of signaling pathways^25–27^. It is likely that metastatic cells gain CNS access through different routes depending on the unique origins and properties of the primary tumor, such as through alterations of choroid plexus tight junctions or via the large arterial vessels in the leptomeningeal space^28,29^. Once CTCs have crossed into the brain parenchyma or leptomeninges, they are challenged to transition from micro-metastases to macro-metastases. This intricate process involves interactions between the CTCs and various components of the neurovascular unit^30–33^. Further, established metastatic lesions co-opt the BBB to become a blood-tumor barrier (BTB) which is characterized by a disorganized vessel structure and leakiness^34^. Heterogeneous permeability of the BBB has been observed in brain metastases of both TNBC and basal-type BC^30^. However, the underlying mechanisms by which BC cells remodel the BBB remain incompletely understood.

Recent studies from our group have revealed that platelet-derived growth factor-B (PDGFB), the most recognized pluripotent angiogenic ligand of the PDGF family^35–37^, and its cognate receptor, platelet-derived growth factor receptor-β (PDGFRβ), play a key role in promoting BCBM^38^. Similarly, Wyss *et al.* demonstrated the importance of BC-derived PDGFA and PDGFB in the formation of brain metastases using murine and human models, supported by patient data showing high *PDGFB* mRNA expression is observed in the primary tumors of patients who developed brain metastasis^39^. Importantly, these pre-clinical findings are supported by increased expression of PDGFB in primary human breast tumors when compared to normal breast tissue^40^. High PDGFB expression also correlated with decreased overall survival (OS) and distant metastasis-free survival (DMFS), as well as shortened time to brain metastatic recurrence^38,40^. Importantly, these studies stopped short of identifying the cellular and molecular mechanisms through which tumor-derived PDGFB promotes CNS metastatic spread, a knowledge gap we address in the present study.

PDGFB is well-known for its role as a pro-angiogenic growth factor in normal physiology^35–37^, where its dominant function is endothelial-to-pericyte paracrine signaling through PDGFRβ to provide vessel integrity and regulate neo-angiogenesis^41,42^. Consequently, it also influences the BBB^43,44^. Previous studies have shown that PDGFB regulates BBB permeability in various pathogenic conditions, such as cocaine-mediated impairment and *Escherichia coli* meningitis infection^45,46^. Additionally, abnormal PDGF signaling has been associated with vascular alterations and BBB dysfunction in glioma^47,48^. In addition, PDGFRβ expression is commonly observed in reactive astrocytes and pericytes^49,50^. Some groups have also reported expression in the microvascular endothelial cells (ECs) forming the BBB^51–54^. Given this established role for PDGFB signaling in angiogenesis, vascular remodeling and BBB biology, we hypothesized that tumor-derived PDGFB alters the CNS vascular niche through interactions with PDGFRβ-positive components of the neurovascular unit in the brain TME, ultimately enhancing metastatic growth in the CNS.

Herein, we present evidence that tumor-derived PDGFB promotes endothelial sprouting, angiogenesis and vascular barrier dysfunction through direct and indirect interactions with cellular populations of the CNS neurovascular unit. Utilizing complementary 2D and 3D *in vitro* models and *in vivo* approaches including a new model for studying spinal metastases and LMD, we found that tumor-derived PDGFB enhances metastatic progression to not only the brain, but also the spinal cord and leptomenigeal space. These pre-clinical findings were validated in patient transcriptomic data suggesting that breast tumor-derived PDGFB drives pro-metastatic angiogenic reprogramming specific for CNS metastatic spread.

## Materials and Methods

### 3D endothelial spheroid and angiogenesis assays

3D HBMEC and HUVEC spheroids were generated using the method previously reported by Shah and colleagues, with slight modifications^55^. Briefly, 1 x 10^3^ cells per 25 μL were suspended via hanging drop method onto the lid of a PBS-filled 15 cm^2^ culture plate and cultured for 24 hours. Spheroids were gathered and reconstituted in basal ECM solution containing 5% FBS along with 40% methylcellulose to hinder settling. The spheroid suspension was mixed with neutralized type I collagen (Corning, #354236), immediately placed on pre-warmed 24-well plates, and allowed to solidify. After 30 minutes, 1 mL of basal ECM, with or without recombinant human platelet-derived growth factor B (rhPDGF-BB, R&D #220-BB-010), recombinant human vascular endothelial growth factor A (rhVEGFA, R&D #293-VE), or 1 mL of CM, was introduced to the gel and incubated for 24 hours. Spheroids were then observed under an EVOS Core XL microscope at a magnification of 20X to monitor the formation of sprouts. Details of quantification and analysis are in Supplemental Methods.

### Microfluidic device fabrication and permeability assay

Microfluidic lumen devices were prepared as previously described, with further detail in Supplemental Methods^56^. After fabrication, lumens were coated overnight at 37°C with 100uL of 100µg/mL fibronectin (Fisher Scientific, #FC010) in Attachment Factor (Cell Systems, #AZ0-201). HBMECs (Cell Systems) were cultured to 85% confluency before harvesting using Passage Reagent Group (Cell Systems, #4Z0-800) and the following morning, 1 x 10^5^ HBMECs were added to each empty lumen. Devices were then placed in humidified dishes and flip steps were completed for 1.5 hours. Non-adhered cells within lumens were removed and fresh media was added and devices were cultured overnight. From Day 1-2, lumen devices were imaged and cultured with normal Lonza EGM-2 media or CM from MB-231-PDGFB/1187-PDGFB and respective controls. Fluorescent tracer (20 kDa FITC-Dextran, Invitrogen #D1820) was resuspended in cell culture media at a concentration of 12.5mg/mL. On Day 2, after clearing and imaging the vessel, 50mL of FITC-dextran solution was introduced into the empty inlet port of the lumen device. Images were taken every 5 seconds for 120 seconds using a 5X objective on a Nikon AX confocal microscope to track vessel permeability. Time-lapse images were then recorded and processed with a custom MATLAB code used to calculate apparent vessel permeability, defined as the net rate of tracer efflux across the endothelial wall into the surrounding ECM^57^.

### Xenograft and Allograft Experimental Metastasis Studies

All animal use was in compliance with University Laboratory Animal Resources regulations under Protocol 2007A0120 (principal investigator: G.M. Sizemore) and approved by the OSU Institutional Animal Care and Use Committee. To investigate widespread metastasis, 1.75 ×10^5^ breast cancer cells were injected into the left ventricle of NSG mice in 200µL phenol red free Opti-MEM^TM^ (ThermoFisher, #11058021)^58^. To investigate LMD, 1.75 ×10^5^ breast cancer cells were injected retro-orbitally in 150µL phenol red free Opti-MEM^TM^. Upon the first mouse reaching early removal criteria (ERC) for each injection model, animals were euthanized via CO_2_ inhalation and perfused with PBS followed by 4% PFA. Additional details are presented in Supplemental Methods.

### Other Methods

Complete and detailed protocols for cell culture, preparation of conditioned media, quantitative real-time PCR (qRT-PCR), western blotting, 3D invasion assays, BBB spheroid development and immunofluorescent imaging, BBB spheroid cell migration, BBB spheroid permeability, cell viability, enzyme-linked immunosorbent assays, immunostaining of cells, frozen tissue, and paraffin tissue, bioluminescent IVIS imaging, *in vivo* Evans blue permeability studies, histopathologic analysis of spinal metastasis, analysis of human breast cancer patient data, and statistical analysis can be accessed in Supplemental Methods. Cell lines with culture conditions are listed in **Supp. Table 1** and antibodies are listed in **Supp. Table 2.**

## Results

### Tumor-derived PDGFB promotes angiogenic responses in brain endothelial cells

To find appropriate experimental tools, we assessed PDGF-BB secretion across BC cell lines and found variable expression and no association with molecular subtype **(Supp. Fig. 1A)**. *PDGFRA* and *PDGFRB* mRNA are minimally expressed in BC lines **(Supp. Fig. 1B-C)** supporting prior studies^59^. We then generated PDGFB-overexpressing variants of the MDA-MB-231 (Basal-B/TN subtype) and HCC1187 (Basal-A/TN subtype) cells^60^, hereafter MB-231-PDGFB and 1187-PDGFB for further study. Successful PDGFB overexpression was confirmed by western blot and ELISA **(Supp Fig. 1D-E)**.

To evaluate potential CNS neurovascular unit responsiveness to PDGF-BB, we evaluated PDGFRβ in primary human astrocytes (HAs), human brain vascular pericytes (HBVPs), and human brain microvascular cells (HBMECs). HBVPs and HAs are known to express PDGFRβ^38,61,62^, and treatment with recombinant human PDGF-BB (rhPDGF-BB) confirmed activation at phospho-tyrosine 1009 in both cell types **(Fig. 1A)**. Uniquely, HBMECs express PDGFRβ, which contrasts with lack of expression in human umbilical vein endothelial cells (HUVECs) **(Fig. 1B)**. Furthermore, we observed high phospho-PDGFRβ in HBMECs upon stimulation with rhPDGF-BB **(Fig. 1B)**. These findings demonstrate that multiple cell types within the CNS neurovascular unit are responsive to PDGF-BB signaling. The observation that HBMECs exhibited functional PDGFRβ activation relative to the non-brain HUVECs, supported an unknown mechanism by which tumor-derived PDGFB can communicate with the CNS vasculature. As such, we focused subsequent mechanistic work on the brain microvascular endothelial cells.

**Figure 1.**
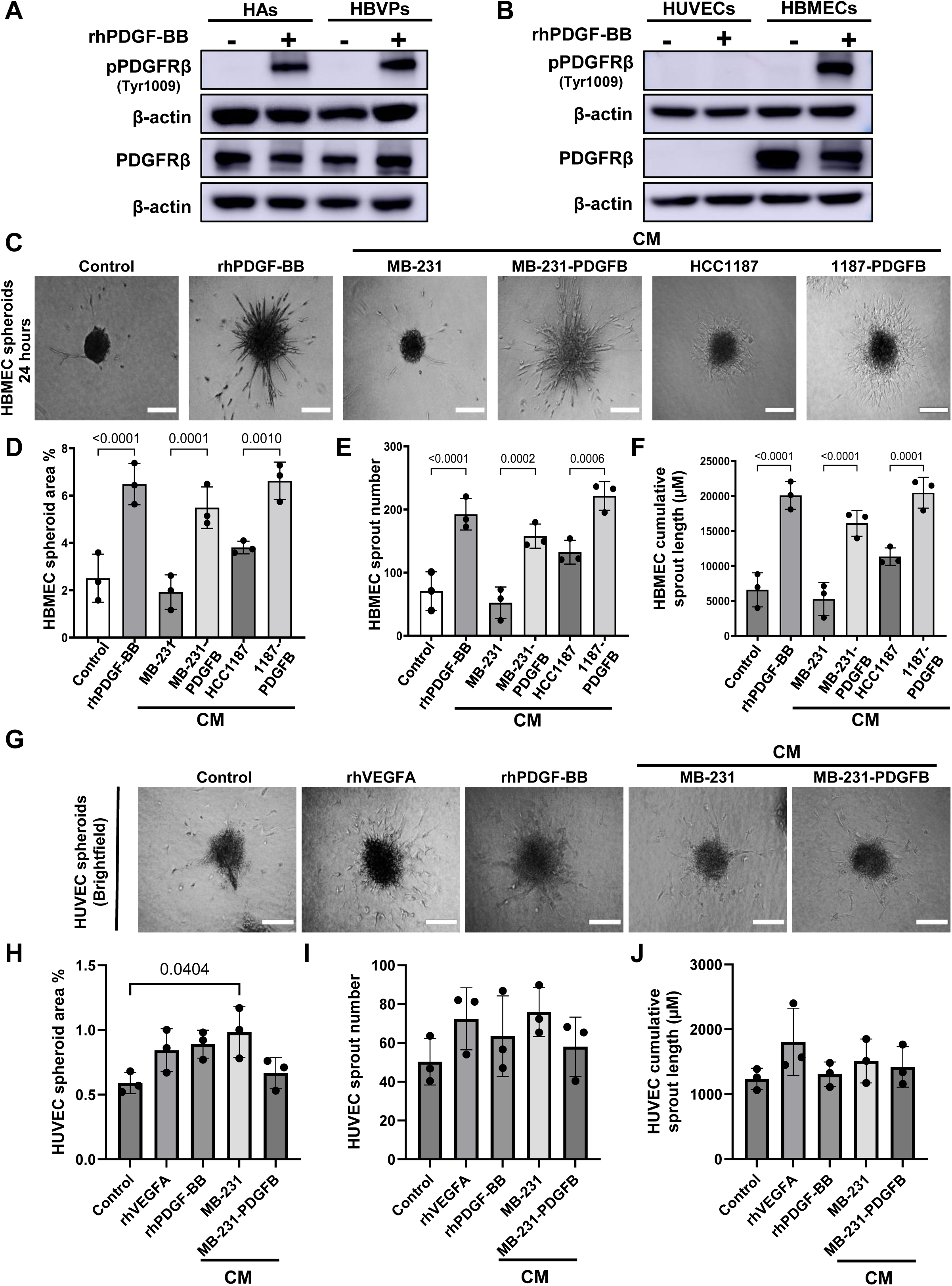
Tumor-derived PDGFB promotes angiogenic responses in brain endothelial cells. **(A)** Western blot analysis of total PDGFRβ and phosphorylated PDGFRβ (Tyr1009) in human astrocytes (HAs) and human brain vascular pericytes (HBVPs) following treatment with rhPDGF-BB (100 ng/mL) for 3 minutes. **(B)** Western blot analysis of total PDGFRβ and phosphorylated PDGFRβ (Tyr1009) in human brain microvascular endothelial cells (HBMECs) and human umbilical vein endothelial cells (HUVECs) following treatment with recombinant human PDGF-BB (100 ng/mL) for 3 minutes. **(C)** Representative images of 3D HBMEC spheroids treated with vehicle control, recombinant human PDGF-BB (100 ng/mL), CM-MB-231, CM-MB-231-PDGFB, CM-HCC1187, or CM-1187-PDGFB for 24 hours. Scale bar, 100 μm. **(D-F)** Quantification of 3D HBMEC (B) spheroid area (%), (C) sprout number, and (D) cumulative sprout length. Data are presented as mean ± SEM from biological replicates (n=3). **(G)** Representative images of 3D HUVEC spheroids treated with vehicle control, VEGFA (50 ng/mL), recombinant human PDGF-BB (100 ng/mL), CM-MB-231, or CM-MB-231-PDGFB for 24 hours. Scale bar, 100 μm. **(H-J)** Quantification of 3D HUVEC (F) spheroid area (%), (G) sprout number, and (H) cumulative sprout length. Data are presented as mean ± SEM from biological replicates (n=3). Alt Text: Western blots of protein expression in various cell types. Microscope images of brain endothelial cell spheroids and graphs showing differences in spheroid sprouting after treatment.

The impact of tumor-derived PDGFB on CNS vascular niche remodeling was first assessed by comparing the angiogenic potential of conditioned media (CM) from MB-231-PDGFB and 1187-PDGFB cells to their respective controls in HBMEC (PDGFRβ-positive) and HUVEC (PDGFRβ-negative) spheroids. Consistent with functional PDGFRβ signaling in HBMECs, rhPDGF-BB treatment and CM from MB-231-PDGFB and 1187-PDGFB cells significantly increased HBMEC spheroid area percentage, sprout number, and cumulative sprout length compared to respective control HBMEC spheroids **(Fig. 1C-F)**. Together, demonstrating that tumor-derived PDGF-BB can promote angiogenic behavior in brain microvascular endothelial cells. To test whether PDGF-BB contributes directly to this phenotype, HBMEC spheroids were treated with neutralizing antibodies targeting each PDGF isoform. While none of the neutralizing antibodies significantly blocked sprouting, the anti-PDGF-BB produced the largest reduction in each measured endpoint **(Supp. Fig. 2A-D)**. In contrast, HUVEC spheroids exhibited a different phenotypic response. Although CM-MB-231 treatment induced a significant increase in HUVEC spheroid area **(Fig. 1G,H)**, no corresponding increase in sprout number or length was detected **(Fig. 1G,I,J)**. Furthermore, CM-MB-231-PDGFB did not significantly alter HUVEC spheroid area, sprout number, or sprout length relative to control **(Fig. 1G-J)**. These findings demonstrate that tumor-derived PDGFB can preferentially promote measurable angiogenic responses in brain microvascular endothelial cells, consistent with a CNS-specific vascular remodeling effect.

To determine whether the increased sprouting observed in the 3D HBMEC spheroids was a result of PDGFB-induced changes in endothelial cell viability, we evaluated HBMEC proliferation in 2D culture. Here, only the addition of FBS significantly altered HBMEC viability **(Supp. Fig. 2E)**. We further assessed whether PDGFB contributed to cell autonomous changes to viability and invasion in the MB-231 and/or HCC1187 variants given detectable expression of *PDGFRB* and *PDGFRA*, respectively in these BC cells **(Supp. Fig. 1A,B)**. Importantly, there were no observed changes in either 2D or 3D growth **(Supp. Fig. 3)**. Combined, these results suggest any observed pro-metastatic effects of tumor-derived PDGFB are likely mediated by CNS vascular remodeling rather than cell-autonomous effects on BC cells.

### Tumor-derived PDGFB disrupts BBB integrity

Given prior evidence implicating PDGFB in BBB regulation^43–48^, we next examined whether tumor-derived PDGFB alters BBB integrity using a previously established *in vitro* 3D tri-culture BBB model^63^. These self-assembling BBB spheroids consist of an HA core followed by HBVPs and then HBMECs form the outer layer of the spheroid **(Supp. Fig. 4)**. In this system, rhPDGF-BB and CM from both MB-231-PDGFB and 1187-PDGFB cells significantly increased cell migration from the BBB spheroid compared to respective controls **(Fig. 2A-B)**. As increased permeability is known to mark BBB dysfunction^64,65^, we examined permeability of high molecular weight dextran into the BBB spheroids. Treatment with rhPDGF-BB and CM from MB-231-PDGFB and 1187-PDGFB cells significantly increased dextran permeability compared to controls **(Fig. 2C-D)**. Overall, these data suggest tumor derived-PDGFB can modulate BBB integrity within multicellular BBB models containing neurovascular HA, HBVPs and HBMEC populations.

**Figure 2.**
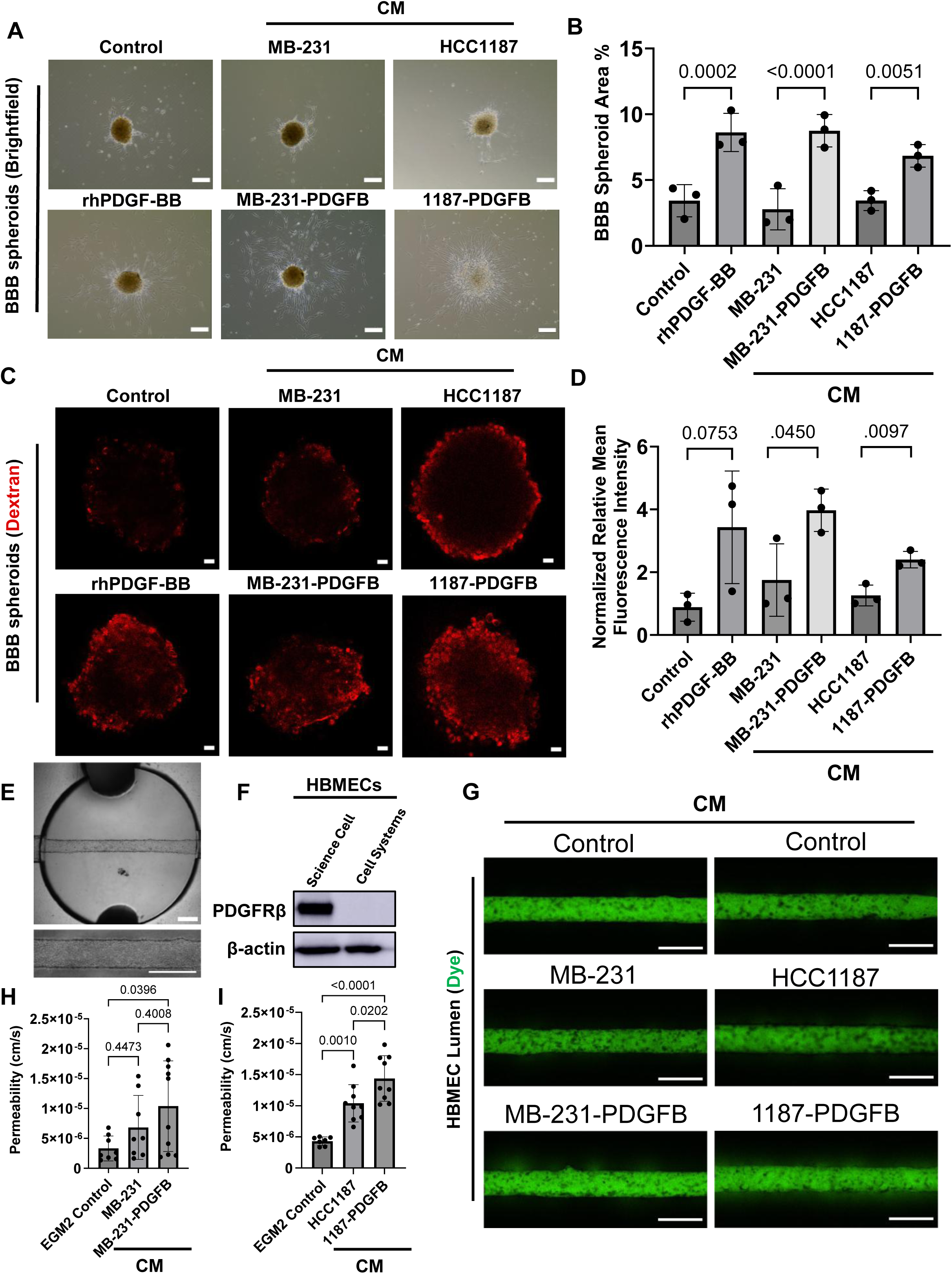
Tumor-derived PDGFB disrupts BBB integrity and increases permeability in BBB *in vitro* model systems. **(A)** Representative brightfield images of tri-culture BBB spheroids composed of HBMECs, HAs, and HBVPs following treatment with vehicle control, rhPDGF-BB (100 ng/mL), CM-MB-231, CM-MB-231-PDGFB, CM-HCC1187, or CM-1187-PDGFB for 24 hours. Scale bar, 100 μm. **(B)** Quantification of migrated area surrounding BBB spheroids in panel A expressed as percentage area covered by migrated cells from the core. Data represents the mean of biological replicates ± SEM (n=3). **(C)** Representative confocal images demonstrating permeability of BBB spheroids to TRITC-dextran (155 kDa; 50 μg/mL). Scale bars, 20 μm. **(D)** Quantification of TRITC-dextran penetration measured at a depth of 70 μm from the spheroid surface and normalized to spheroid area. **(E)** Representative bright field images of engineered HBMEC microvessel devices. Scale bar, 500 μm. **(F)** Western blot analysis of PDGFRβ protein expression in HBMEC cells from Science Cell or Cell Systems vendors. Cell Systems HBMECs were utilized in the engineered microvessel experiments. **(G)** Representative fluorescent images of engineered HBMEC microvessels following treatment with control media, CM-MB-231, CM-MB-231-PDGFB, CM-HCC1187, or CM-1187-PDGFB. Scale bars, 500 μm. **(H-I)** Quantification of apparent permeability in microvessels treated with (H) MB-231 and (I) HCC1187-derived CM. Data are presented as mean ± SEM; control (n=9), CM-MB-231 (n=8), CM-MB-231-PDGFB (n=10), CM-HCC1187 (n=9), and CM-1187-PDGFB (n=10). Alt Text: Brightfield and fluorescent images of BBB spheroids and engineered brain endothelial cell lumens and graphs showing quantification of spheroid are and permeability to dye.

To determine whether PDGFB-mediated barrier dysfunction extends beyond PDGFRβ-positive endothelial systems, we utilized an engineered microvessel platform **(Fig. 2E)** generated from a brain endothelial cell population lacking PDGFRβ expression **(Fig. 2F)**. Microvessels treated with CM-MB-231-PDGFB had significantly increased apparent permeability as tracked by Dextran dye compared to control vessels **(Fig. 2G,H)**, while microvessels treated with CM-1187-PDGFB had significantly higher apparent permeability than both control and CM-1187 vessels **(Fig. 2G,I).** These findings suggest that PDGFB-mediated vascular dysfunction can also occur in endothelial systems lacking detectable PDGFRβ expression, supporting the existence of direct and indirect mechanisms of CNS vascular remodeling.

### PDGFB induces endothelial tight-junction remodeling

Given the direct responsiveness of PDGFRβ-positive HBMECs to PDGFB, we next examined whether PDGFB could induce alterations in tight junction protein expression given that they are critical for maintaining BBB integrity, and their disruption can lead to increased permeability^66^. Western blot analysis showed a significant decrease in the expression of ZO-1, occludin, and claudin-5 in HBMECs treated with CM-MB-231-PDGFB compared to those treated with control CM **(Fig. 3A-B)**. Similarly, both ZO-1 and occludin were significantly decreased by immunofluorescence in HBMECs treated with CM-MB-231-PDGFB compared to control **(Fig. 3C-F)**. Notably, ZO-1 was mainly observed as solid lines at the periphery of HBMECs treated with CM-MB-231. This distribution became irregular and discontinuous upon treatment with CM-MB-231-PDGFB **(Fig. 3C)**. These findings provide further evidence that tumor-derived PDGFB can directly alter HBMEC biology in a pro-metastatic manner.

**Figure 3.**
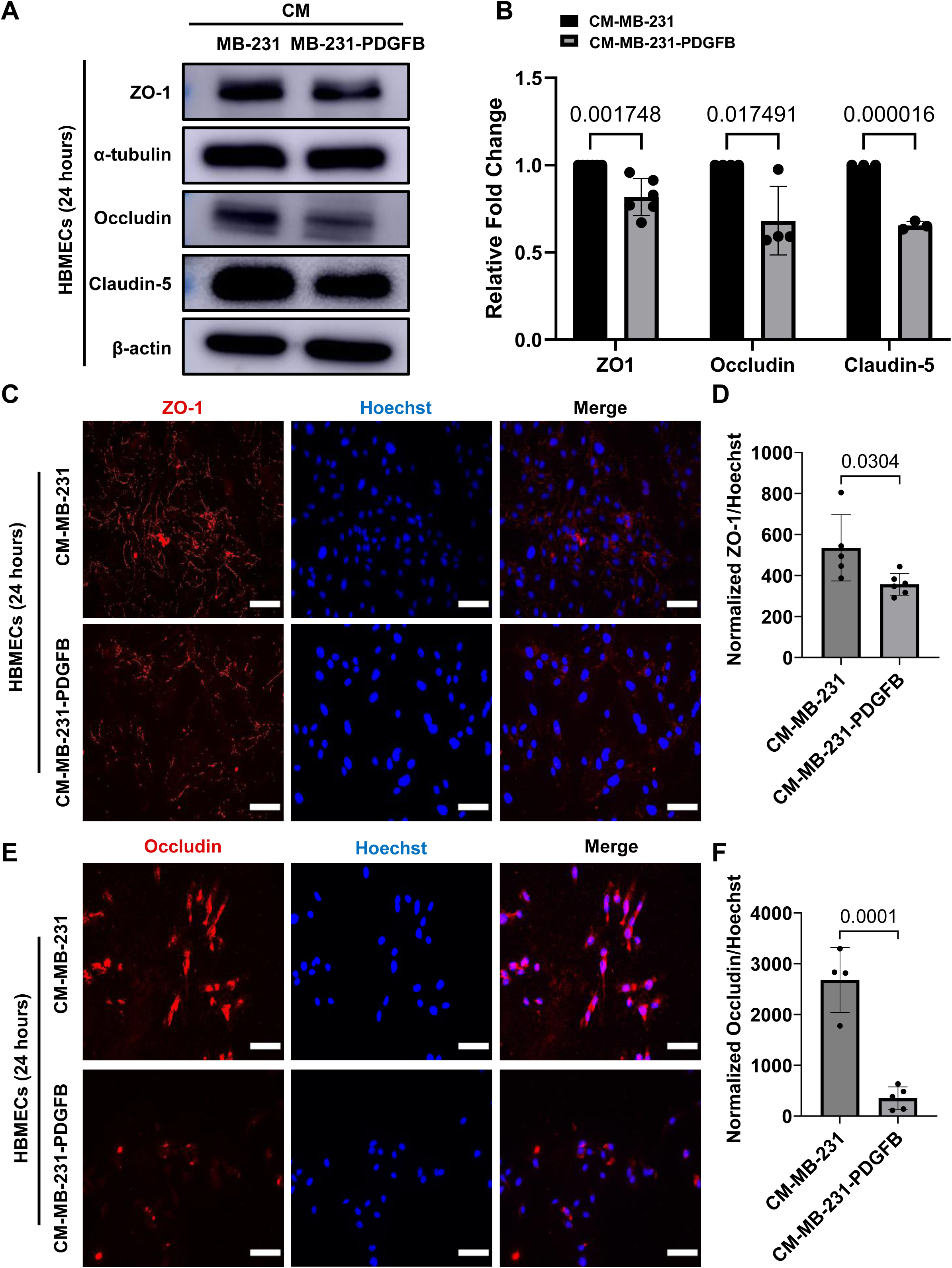
Tumor-derived PDGFB induces loss of endothelial tight-junction proteins. **(A)** Western blot analysis of ZO-1, Occludin, and Claudin-5 expression in HBMECs following treatment with CM-MB-231 or CM-MB-231-PDGFB for 24 hours. Representative blots from independent biological experiments are shown. **(B)** Quantification of protein expression normalized to β-actin and expressed relative to CM-MB-231-treated controls. Data are presented as mean ± SEM from biological replicates (n=3-6). **(C)** Representative immunofluorescence images of ZO-1 localization in HBMEC monolayers following 24-hour treatment with conditioned media. Nuclei were counterstained with Hoechst. **(D)** Quantification of ZO-1 fluorescence intensity normalized to nuclear staining. Data are presented as mean ± SEM (n=4-5). Scale bars, 100 μm. **(E)** Representative immunofluorescence images of Occludin localization in HBMEC monolayers following 24-hour treatment with conditioned media. Nuclei were counterstained with Hoechst. **(F)** Occludin fluorescence intensity normalized to nuclear staining. Data are presented as mean ± SEM (n=4-5). Scale bars, 100 μm. Alt Text: Western blot and fluorescent images of tight junction proteins in treated endothelial cells and graphs showing quantification of tight junction proteins.

### PDGFB promotes CNS metastasis and vascular dysfunction in intracardiac xenograft models

Previous studies by our group have demonstrated a role for tumor-derived PDGFB in intracranial BC tumor growth and brain-specific experimental metastasis following tail vein injection^38^. Thus, we sought to determine whether PDGFB-driven vascular remodeling increases CNS metastatic growth *in vivo*. To this end, luciferase expressing PDGFB overexpressing lines and respective control cells were injected intracardially into immunodeficient NSG mice. Mice injected with either MB-231 control or 231-PDGFB cells exhibited IVIS signal throughout the body including in the head, lung and liver at ERC **(Fig. 4A,B)**. IVIS quantification of the total luminescence flux of the head as well as the body suggested a decrease in brain metastatic burden of the MB-231-PDGFB cells **(Fig. 4C,D)**; however, we consistently noted neurological symptoms in these mice that also could not be confirmed by H&E imaging of the brain (data not shown). Thus, we further investigated the brain-specific metastatic burden of the MB-231 mice by MRI at ERC **(Fig. 4E)**. Interestingly, we observed little to no parenchymal tumor burden in either group; however, we did observe significant LMD in both MB-231-PDGFB and control cell injected mice, with MB-231-PDGFB injected mice exhibiting a more severe LMD burden **(Fig. 4E,F)**. Mice injected with 1187-PDGFB cells displayed significant increases in both total body and cranial metastasis (total head) compared to the control group as measured by IVIS at ERC **(Fig. 4G-I)**. Unlike in the 231 cohort, an increase in parenchymal tumors was easily observed and confirmed by H&E staining **(Fig. 4J)**. Altogether, revealing that increased PDGFB promotes CNS parenchymal metastasis and LMD in xenograft models of BC.

**Figure 4.**
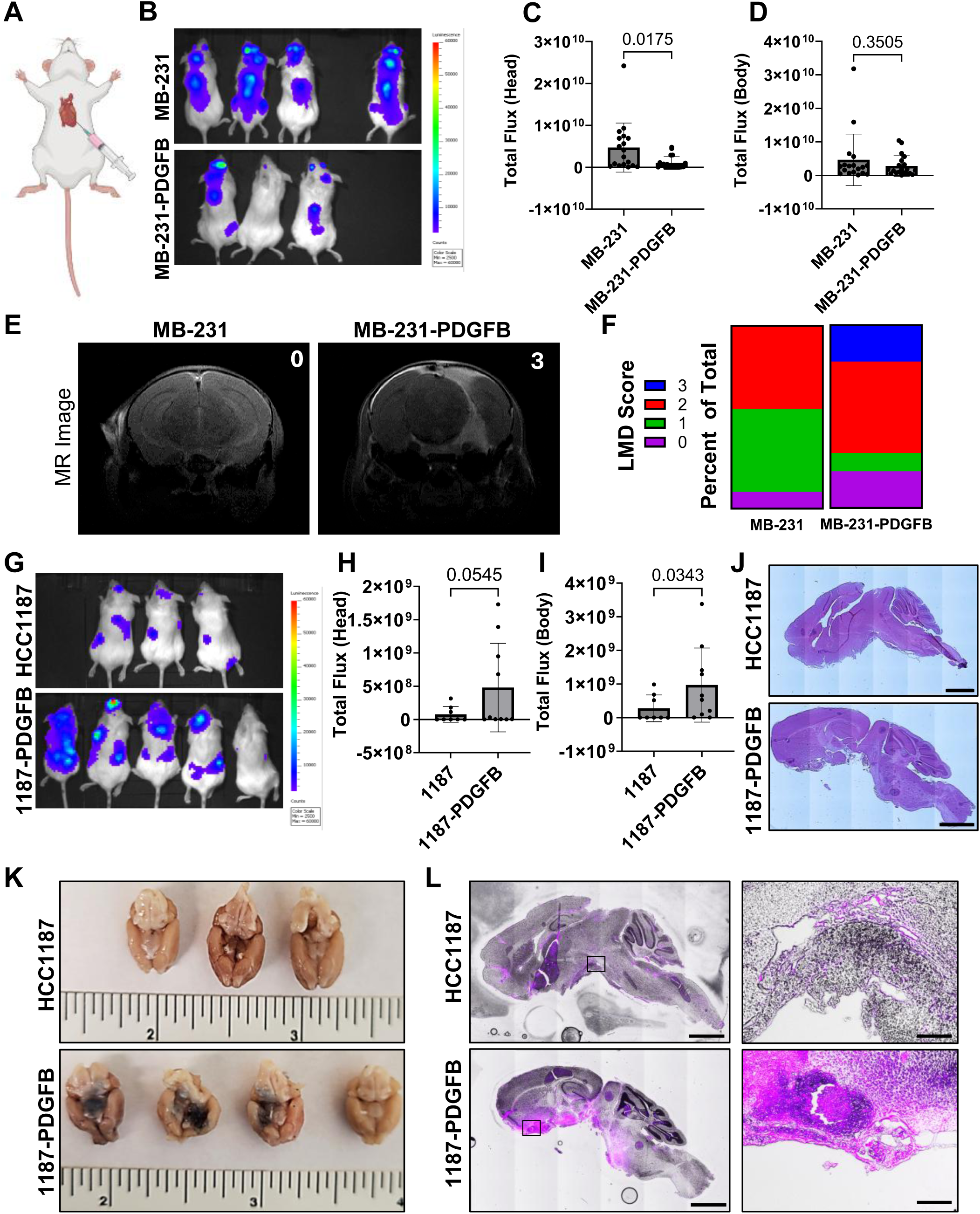
PDGFB promotes CNS metastasis and vascular dysfunction in intracardiac xenograft models. **(A)** Schematic of the intracardiac injection model. **(B)** Representative IVIS images of NSG mice following intracardiac injection of luciferase-expressing MB-231 or MB-231-PDGFB cells. **(C-D)** Quantification of bioluminescent signal in the (C) head and (D) body regions. Data are presented as mean total flux ± SEM. MB-231 (n=18); MB-231-PDGFB (n=20). **(E)** Representative MR images demonstrating leptomeningeal metastatic burden in MB-231 and MB-231-PDGFB cohorts with the LMD severity score in the upper right. **(F)** Radiology scoring of LMD severity. MB-231 (n=11); MB-231-PDGFB (n=10). **(G)** Representative IVIS images of NSG mice following intracardiac injection of luciferase-expressing HCC1187 or 1187-PDGFB cells. **(H-I)** Quantification of bioluminescent signal in the (H) head and (I) body regions. Data are presented as mean total flux ± SEM. HCC1187 (n=8); 1187-PDGFB (n=10). **(J)** Representative H&E images of NSG mice following intracardiac injection of luciferase-expressing HCC1187 or 1187-PDGFB cells. Scale bar, 2000 μm. **(K)** Representative bright field imaging of Evans blue dye penetration in brains of mice injected with HCC1187 (n = 3) or 1187-PDGFB (n = 4) cells. Ruler scale is inches. **(L)** Representative Evans blue dye fluorescence demonstrating vascular permeability within the metastatic lesions formed in the brains of mice injected with HCC1187 or 1187-PDGFB cells. Scale bars, 2000 μm (left) and 200 μm (right). Alt Text: Luminescence, MRI, and histological images showing tumor burden in mice intracardially injected with breast cancer cells.

To validate our *in vitro* findings of PDGFB-mediated BBB modulation, we further assessed *in vivo* BBB permeability in the HCC1187 intracardiac model using Evans blue dye, which can only enter the CNS when the BBB/BCSFB is compromised. Evans blue dye was administered intraperitoneally at ERC, and post-mortem analysis of the brains revealed an increase in the uptake of Evans blue dye in the brain metastases of mice injected with 1187-PDGFB cells compared to control cells **(Fig. 4K,L)**. Therefore, increased BC-derived PDGFB can alter BBB/BCSFB function *in vivo* suggesting paracrine dysregulation of the BBB/BCSFB from the injected BC cells.

### PDGFB enhances spinal and leptomeningeal metastatic colonization

To further investigate PDGFB’s role in LMD, we performed retro-orbital (RO) injections of luciferase expressing PDGFB overexpressing cells and respective control cells, which more precisely delivers cells into the CNS and reduces the extra-cranial metastatic burden (e.g., lung outgrowth) that led to early removal **(Fig. 5A)**. In mice injected with MB-231-PDGFB cells, we again observed a significant reduction in head luminescence signal by IVIS at ERC **(Fig. 5B,C)** matching the result observed with intracardiac injection **(Fig. 4B,C)**. However, this result was concomitant with a significant increase in spine luminescence signal **(Fig. 5B,D)**. To confirm this signal was indeed due to metastases in the spines of the mice, representative sections of cervical, thoracic, and lumbar spinal cord and cauda equina were examined by a board-certified comparative pathologist. Indeed, mice injected with MB-231-PDGFB cells had multiple spinal masses in distal aspects of the spinal cord and cauda equina (n = 4/4 and 3/5 in respective cohorts) while mice injected with MB-231 cells did not (n = 0/6) **(Fig. 5E)**. Further histologic examination confirmed metastatic tumor infiltration of spinal nerves, dorsal root ganglia, leptomeninges, and spinal cord structures **(Fig. 5F)**. In mice injected with 1187-PDGFB cells, we observed no change to head or spinal IVIS signal at ERC **(Fig. 5G-I)**. However, we found a significant increase in spinal metastases in the 1187-PDGFB group compared to control by quantification of anti-human nucleolin immunostaining throughout the spinal columns of these mice **(Fig. 5J,K)**. These studies strongly support a pro-metastatic role for BC-derived PDGFB in the development of LMD and reveal a technical limitation in using IVIS as an imaging modality for LMD and spinal metastases.

**Figure 5.**
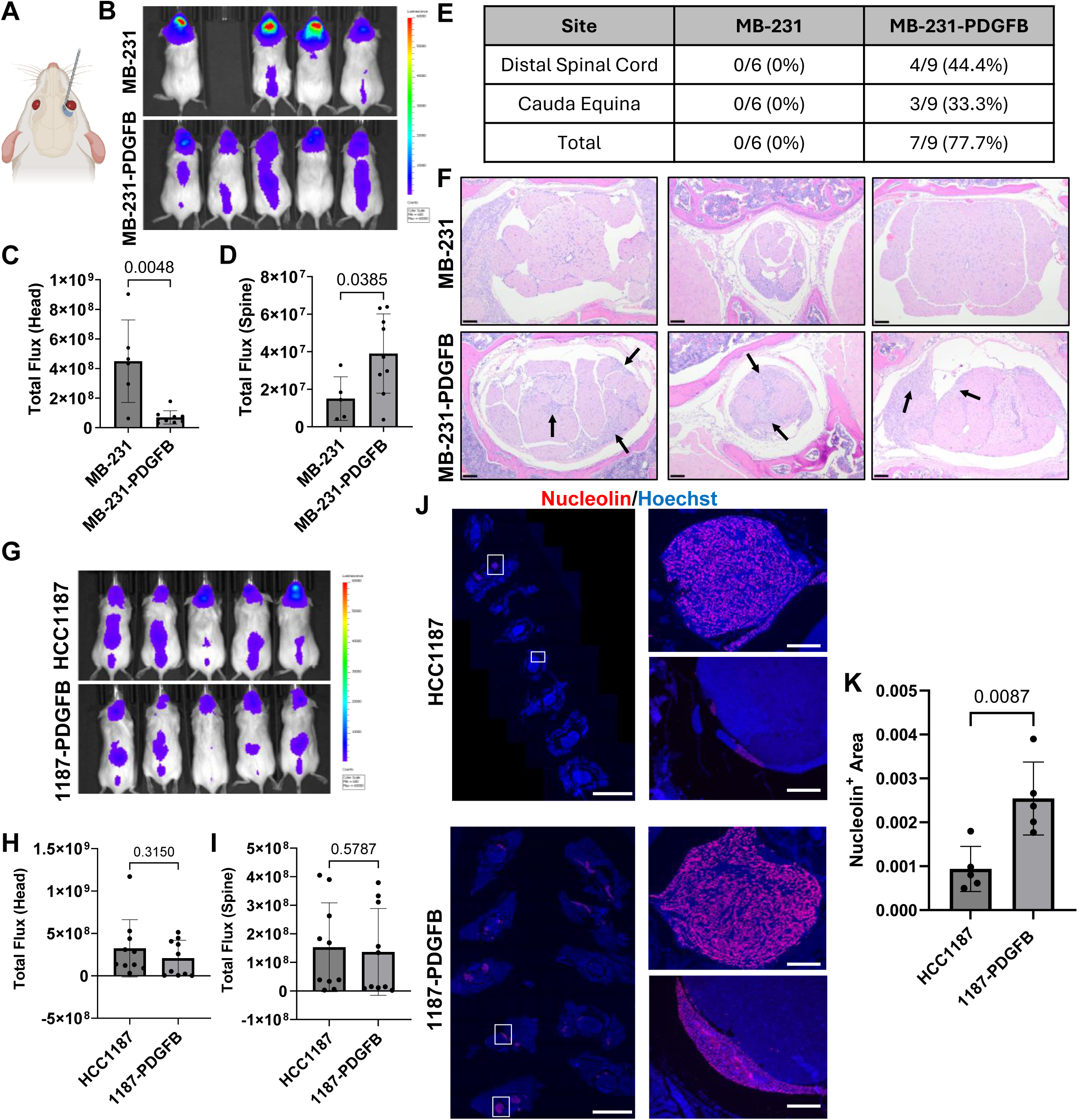
PDGFB enhances spinal and leptomeningeal metastatic colonization. **(A)** Schematic of the retro-orbital injection model. **(B)** Representative IVIS images of NSG mice following retro-orbital injection of luciferase-expressing MB-231 or MB-231-PDGFB cells. **(C-D)** Quantification of bioluminescent signal in the (C) head and (D) spine. Data are presented as mean total flux ± SEM. MB-231 (n=6); MB-231-PDGFB (n=9). **(E)** Frequency of metastatic involvement at distal spinal cord and cauda equina sites. **(F)** Representative H&E images of spinal metastases. Arrows indicate infiltration of neoplastic cells into spinal nerves, meninges, and spinal cord structures. Scale bar, 100 μm. Total magnification, 100X. **(G)** Representative IVIS images following retro-orbital injection of HCC1187 or 1187-PDGFB cells. **(H-I)** Quantification of bioluminescent signal in the (H) head and (I) body. Data are presented as mean total flux ± SEM. HCC1187 (n=10); HCC1187-PDGFB (n=10). **(J-K)** Representative (J) immunofluorescence images and (K) quantification of anti-human nucleolin marking spinal metastatic burden. Scale bars, 5000 μm (left) and 200 μm (right). Alt Text: Luminescence, fluorescent, and histological images showing tumor burden in mice retro-orbitally injected with breast cancer cells.

### PDGFB promotes CNS vascular remodeling and metastatic growth in an immunocompetent model of breast cancer

Given the importance of the immune system in metastasis, we sought to confirm the observed vascular modulating role of PDGFB in a syngeneic mouse model with an intact immune system. To this end, we performed intracardiac injections in wild-type FVB/N mice of the murine mammary tumor DB7 cell line with or without shRNA mediated silencing of *Pdgfb*^38^ **(Fig. 6A)**. Brains were harvested when the first mouse met ERC and tumor burden was evaluated by H&E **(Fig. 6B)**. All animals injected with DB7 shCtrl (n=4) developed overt brain metastases, whereas only 2/4 injected with DB7 sh*Pdgfb* developed any observable brain metastases. When quantified, the overall tumor burden **(Fig. 6C)** and number of tumors per brain slice **(Fig. 6D)** were significantly decreased in the DB7 sh*Pdgfb* group. We then stained brain slices for CD31 to investigate differences in brain tumoral vasculature and found intratumoral CD31 positivity was significantly decreased upon PDGFB silencing (**Fig. 6E-F**).

**Figure 6.**
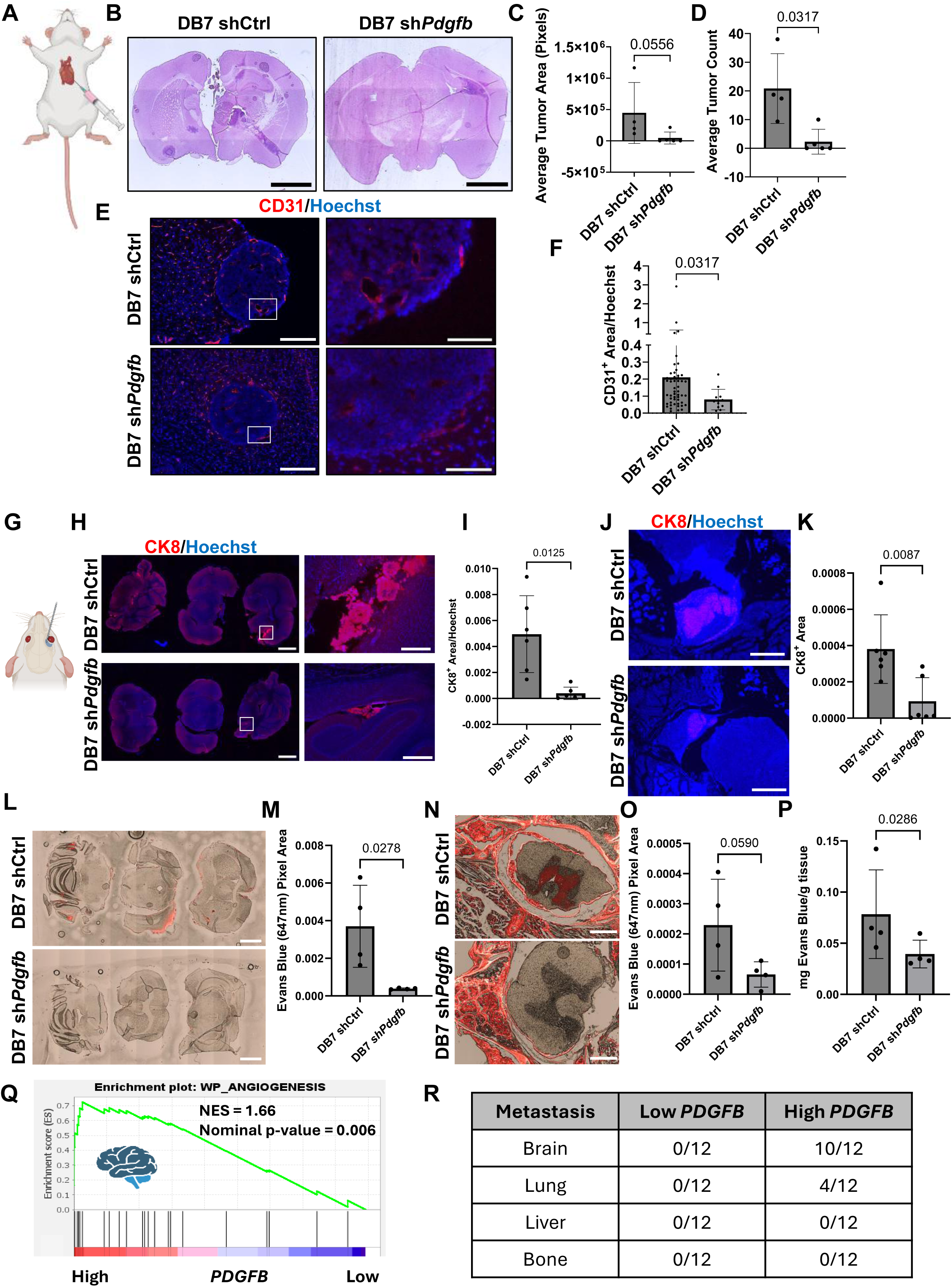
PDGFB promotes CNS vascular remodeling and metastatic growth in an immunocompetent model of breast cancer. **(A)** Schematic of the intracardiac injection model. **(B)** Representative H&E images of brain metastases following intracardiac injection of DB7 shCtrl or DB7 sh*Pdgfb* cells into FVB/N mice. Scale bar, 2000 μm. **(C-D)** Quantification of (C) average tumor area and (D) tumor number per section. Data are presented as mean ± SEM. shCtrl (n=4); sh*Pdgfb* (n=5). **(E)** Representative immunofluorescence images of CD31-positive tumor vasculature. Scale bars, 2000 μm (left) and 100 μm (right). **(F)** Quantification of CD31-positive area normalized to Hoechst. Data are presented as mean ± SEM. **(G)** Schematic of the retro-orbital injection model. **(H)** Representative immunofluorescence images of cytokeratin-8 (CK8)-positive metastatic brain tumor burden following retro-orbital injection of DB7 shCtrl or DB7 sh*Pdgfb* cells. Scale bars, 2000 μm (left) and 275 μm (right). **(I)** Quantification of CK8-positive brain metastatic tumor area normalized to Hoechst. Data are presented as mean ± SEM. shCTRL (n=6); sh*Pdgfb* (n=6). **(J)** Representative immunofluorescence images of CK8-positive metastatic spine tumor burden following retro-orbital injection of DB7 shCtrl or DB7 sh*Pdgfb* cells. Scale bars, 500 μm. **(K)** Quantification of CK8-positive brain metastatic tumor area normalized to Hoechst. Data are presented as mean ± SEM. shCtrl (n=6); sh*Pdgfb* (n=6). **(L)** Representative Evans blue dye fluorescence (647nm) demonstrating vascular permeability within metastatic brain lesions from mice injected with DB7 shCtrl or DB7 sh*Pdgfb* cells. Scale bars, 2000 μm. **(M)** Quantification of brain Evans blue dye in L. Data are presented as mean ± SEM. shCtrl (n=4); sh*Pdgfb* (n=4). **(N)** Representative Evans blue dye fluorescence demonstrating vascular permeability within metastatic spine lesions from mice injected with DB7 shCtrl or DB7 sh*Pdgfb* cells. Scale bars, 200 μm. **(O-P)** Quantification of spine Evans blue dye by (O) pixel area and (P) mg/tissue (right). Data are presented as mean ± SEM. shCTRL (n=4); sh*Pdgfb* (n=4). **(Q)** Representative enrichment plot (WP_ANGIOGENESIS) demonstrating significant enrichment in patients who developed brain metastases and exhibited high primary tumor PDGFB expression (n=66). **(R)** Summary of significantly enriched angiogenic gene sets across metastatic sites. Gene sets were considered significantly enriched when FDR q-value < 0.25.Alt Text: Fluorescent and histological images showing tumor burden in mice intracardially or retro-orbitally injected with breast cancer cells. Graph and table showing angiogenic gene set markers in human breast cancer patients with high and low *PDGFB*.

To evaluate LMD and spinal metastases in this model system, we also performed RO injections of DB7 shCtrl and sh*Pdgfb* cells **(Fig. 6G)**. Consistent with the intracardiac studies, loss of PDGFB significantly reduced brain **(Fig. 6H-I)** and spinal metastatic **(Fig. 6J-K)** burden, as assessed by cytokeratin (CK8) immunostaining at ERC. These results point to tumor derived PDGFB as a strong driver of CNS metastasis, independent of the immune system. To confirm that tumor derived PDGFB alters permeability of spinal metastases, we again performed RO injections of the DB7 shCtrl and sh*Pdgfb* cell lines in FVB/N mice. Evans blue dye was administered as above at ERC. Histological evaluation of the Evans blue post-mortem revealed that uptake of the dye was significantly diminished in the brains of mice injected with DB7 sh*Pdgfb* cells **(Fig. 6L-M)**. Additionally, we found that Evans blue uptake was decreased in the spinal cord and leptomeninges of mice injected with DB7 sh*Pdgfb* by both histology **(Fig. 6N-O)** and quantification of Evans blue signal extracted from spines via *N,N*-Dimethylformamide **(Fig. 6P)**. Collectively, these findings demonstrate that tumor-derived PDGFB promotes CNS vascular remodeling, barrier dysfunction, and metastatic outgrowth in both xenograft and immunocompetent models.

### Angiogenic gene signatures are enriched in patients with high PDGFB tumors that develop brain metastases

To determine the association between primary breast tumor *PDGFB* mRNA expression and angiogenesis-related pathways in human patients that develop BC-CNS metastases, we performed GSEA on patients with known metastatic recurrence in the METABRIC BC patient cohort^67,68^. Primary tumor transcriptomes were stratified by median *PDGFB* mRNA expression into high versus low *PDGFB* groups and analyzed independently across patients who developed brain, lung, liver, or bone metastasis **(Supp. Fig. 5A)**. In an initial unsupervised analysis, WP_ANGIOGENESIS was among the top 20 significantly enriched gene sets of patients with high primary tumor *PDGFB* expression and brain metastatic recurrence (nominal *P* = 0.006, NES = 1.66, **Fig. 6Q**). This finding prompted us to perform a supervised analysis restricted to angiogenesis-related gene sets across each metastatic group. We found that 10 of 12 and 4 of 12 angiogenesis-related gene sets were significantly enriched in high *PDGFB* tumors from patients with brain and lung metastasis, respectively **(Supp. Table 3, Fig. 6R, Supp. Fig. 5B).** In contrast, no angiogenesis-related gene sets were significantly enriched in high *PDGFB* tumors from patients with liver or bone metastasis **(Fig. 6R, Supp. Fig. 5C-E).** Together, these findings provide clinical support for a PDGFB-associated vascular remodeling program in patients who subsequently develop brain metastases.

## Discussion

PDGFB is well known to promote BC progression through multiple context-dependent autocrine and paracrine mechanisms^59^. We have previously shown that PDGFB-to-PDGFRβ signaling preferentially promotes metastasis to the brain^38^. However, the cellular-molecular mechanisms underlying this CNS tropism remained poorly understood. Here, we provide evidence that tumor-derived PDGFB remodels the CNS vascular niche through direct and indirect interactions with the brain endothelium, leading to enhanced endothelial activation, angiogenesis, and vascular barrier dysfunction. These vascular changes were associated with increased metastatic colonization of the brain parenchyma, leptomeningeal space, and spinal cord across multiple experimental models. Collectively, our findings support a model in which tumor-derived PDGFB promotes BC-CNS progression by reshaping the CNS vascular microenvironment to create a permissive niche for metastatic growth.

When studying the neurovascular unit, it is notable that multiple cell types express PDGFRβ. This included the HBMECs, which express high levels of PDGFRβ compared to the HUVEC model. Although PDGFRs are not typically expressed in endothelial cell populations, this observation supports a study conducted by Smits *et al*., which demonstrated that the microvessel ECs comprising the BBB express functional PDGFRβ whereas the ECs that form large cerebral blood vessels do not^54^. Interestingly, CM from PDGFB overexpressing cells significantly increased the sprouting angiogenic potential of HBMECs in a PDGFB-dependent manner while this effect was not observed in HUVEC spheroids. We further assessed the ability of tumor-derived PDGFB to modulate 3D BBB spheroids consisting of HBMECs, HAs, and HBVPs^69,70^. CM from PDGFB overexpressing cells significantly increased BBB spheroid area and increased permeability as measured by dextran penetrance. To dive deeper mechanistically, we employed a static 3D engineered HBMEC microvessel to more accurately model the reported phenotype of large brain vessel endothelial cells. Even without apparent PDGFRβ, tumor-derived PDGFB significantly increased permeability of engineered vessels to fluorescent dye compared to control, supporting a role for PDGFB in CNS vascular remodeling through both direct and indirect vascular mechanisms. Together, these *in vitro* model systems provide complementary approaches for mechanistically investigating BBB biology.

Given the observed brain-specific endothelial PDGFRβ expression, we sought to interrogate the effect on other features of brain endothelial cells. Tight junction proteins are a unique and integral component of an intact BBB^19–21^. To determine if PDGFB could induce changes in tight junction protein expression, HBMECs were treated with CM from MB-231-PDGFB and control cells. Tumor-derived PDGFB significantly reduced protein expression of ZO-1, Occludin, and Claudin-5, identifying a mechanism by which PDGFB-expressing cells can modulate BBB permeability. While previous studies have shown that tumors with high PDGFB expression elevate vascular permeability due to pericyte loss^71^, our study identifies a mechanism by which tumor-derived PDGFB enhances vessel permeability and angiogenesis through direct effects on brain endothelial cells. While *in vitro* BBB models and the 3D HBMEC spheroid angiogenesis model are valuable tools, it’s important to recognize that they do not fully replicate the complex microenvironment of the brain. Thus, we performed confirmatory *in vivo* studies in which intracardiac injection of BC cells showed that PDGFB overexpression increases parenchymal brain metastasis in the HCC1187 cells and LMD in MB-231 cells. Using this injection technique, we also confirmed the *in vitro* permeability findings by Evans blue dye penetration *in vivo*.

Among the most clinically relevant findings of this study, PDGFB markedly enhanced metastatic colonization of both the spinal cord and leptomeningeal space, two understudied complications of BC-CNS progression associated with particularly poor patient outcomes. Here, we supplemented the traditional intracardiac injection approach with a retro-orbital (RO) injection technique, which allows tumor cells to directly enter the brain circulation and for direct drainage of cells into the cavernous sinous and basilar venous plexus, where cells can enter the CSF circulatory system^72,73^. Consistent with this route of delivery, we routinely see leptomeningeal, ventricular, and spinal tumor burden upon injection with each cell line. This suggests that the route by which CTCs enter the CNS is an important step in determining the phenotype of the tumor, and thus more research is required to elucidate these various mechanisms of entry. To our knowledge, we are the first to demonstrate the efficacy of the RO injection for study of LMD. The current models for studying LMD involve invasive and precise stereotactic procedures to deliver cells into the cisterna magna, which requires specialized equipment and bypasses vessel extravasation^74^. The RO injection is simple and can be performed with minimal training and equipment, equipping more researchers to study this devastating form of metastasis. We found that PDGFB overexpression conferred a striking advantage in the ability of all three cell lines tested to successfully colonize the spinal cord parenchyma and leptomeninges in both immune competent and deficient systems. These findings support the hypothesis that PDGFB-mediated vascular remodeling may facilitate metastatic outgrowth in otherwise restrictive CNS compartments. This mechanism is corroborated by the increased Evans blue permeability in the brains and spinal cords of animals injected with high PDGFB expressing BC cells. Our findings are particularly relevant in LMD, as increased permeability of the BCSFB has been shown to be crucial for metastatic outgrowth in this space^75,76^. Ultimately, we hope that this model will facilitate mechanistic evaluation of how CTCs colonize the spinal cord and leptomeningeal space and enable development of new therapeutic strategies for this devastating disease.

Of note, tracking LMD using conventional monitoring by luciferase labeling on cancer cells and *in vivo* imaging by IVIS was challenging. IVIS requires the luciferin substrate to reach tumor cells and we found that efficient biodistribution was lacking in the leptomeningeal space. Given our findings that PDGFB alters vascular permeability, technical difficulties observing signal in CNS lesions are not unsurprising. Consequently, we required the use of MRI to accurately track tumor progression, which highlights the unique difficulties of studying LMD. Moving forward, we provide a practical workflow utilizing RO injections followed by MRI and/or systemic histopathology to directly study LMD.

Importantly, bioinformatic evaluation of the METABRIC cohort^67^ reinforces our experimental observations, demonstrating that high *PDGFB* mRNA expression in the primary tumor is associated with angiogenesis-associated transcriptional programs in patients who subsequently develop brain metastases. This association is limited in patients that developed lung metastases and is absent in patients that went on to develop liver or bone metastases. Importantly, these findings align with and extend our experimental results, providing clinical and transcriptomic evidence linking primary tumor *PDGFB* expression to angiogenesis specifically in BCBM. Presently, data from patients with LMD are sparse, but future work is warranted to determine if *PDGFB* expression plays a prognostic role in predicting which BC patients may have increased risk for developing LMD and if it predicts outcome in patients with established BC-CNS metastasis.

In summary, our work has identified tumor-derived PDGFB as a regulator of the CNS vascular niche during breast cancer progression. PDGFB directly induces vascular changes including angiogenic endothelial cell sprouting and increased BBB permeability through modulation of tight junction protein expression. Tumors with high levels of PDGFB perturb CNS barriers and vascularization *in vivo*, leading to increased metastatic colonization of the brain, spinal cord and leptomeningeal space. Additionally, we establish a practice experimental workflow for studying spinal metastases and LMD in mice, which could provide new opportunities to investigate the unique biology behind these devastating complications of BC-CNS disease.

## Declarations

### Ethics

All animal use was in compliance with University Laboratory Animal Resources regulations under Protocol 2007A0120 (principal investigator: G.M. Sizemore) and approved by The Ohio State University Institutional Animal Care and Use Committee.

### Funding

This study was supported by the National Cancer Institute at the National Institutes of Health (1K22CA218472-01 and R37CA282660, G.M.S.), the Department of War Breast Cancer Research Program (W81XWH-20-1-0047, S.T.S., W81XWH-21-1-0021, G.M.S., and HT9425-25-1-0081, S.T.S.), the National Institute of General Medical Sciences (P20GM121322, P.R.L), and the National Institute of Neurological Disorders and Stroke (R01NS118037, J.P.G). Research reported in this publication was supported by The Ohio State University Comprehensive Cancer Center and the National Institutes of Health (P30CA016058). We specifically thank the Comparative Pathology & Digital Imaging Shared Resource and the Small Animal Imaging Shared Resource for histopathology and mouse imaging support, respectively. The work was further supported by The Ohio State University Comprehensive Cancer Center Pelotonia Intramural Research Program (G.M.S.) and the Pelotonia Scholars Program (S.S.). Any opinions, findings, and conclusions in this material are those of the author(s) and do not necessarily reflect those of the Pelotonia Scholars Program or The Ohio State University.

### Conflict of Interest

The authors declare no potential conflicts of interest.

### Authorship Contributions

**Conceptualization:** N.S.G., J.J.R., S.S., J.J.A., K.A.T., J.W.S., S.T.S., G.M.S. **Data curation:** N.S.G.., J.J.R., S.S., J.J.A., R.L.P., P.A.K., S.S.A., J.M.S., T.A.A., J.W.C., A.A.M., J.G.S., S.A.S., A.J.D., D.S.R., L.K., S.B., K.N.C. **Formal analysis:** N.S.G., J.J.R., S.S., J.J.A., P.L., P.A.K., S.S.A., S.B., K.N.C., J.W.S., S.T.S., G.M.S. **Investigation:** N.S.G., J.J.R., S.S., S.B., K.N.C. **Methodology:** N.S.G., J.J.R., S.S., J.J.A., P.L., R.L.P., Y.R., K.A.T., P.A.K., S.S.A., J.M.S., T.A.A., S.A.S., D.S.R., L.K., M.V., J.P.G., P.R.L., S.B., K.N.C., J.W.S., S.T.S., G.M.S. **Validation:** N.S.G., J.J.R., S.S., J.J.A., P.L., R.L.P., Y.R., S.S.A., J.M.S., S.A.S., S.B., K.N.C., J.W.S. **Visualization:** N.S.G., J.J.R., S.S., J.J.A., P.A.K., S.S.A., K.N.C., J.W.S., G.M.S. **Resources:** M.V., P.R.L., J.W.S., S.T.S., G.M.S., **Supervision:** M.V., J.P.G., P.R.L., J.O.W., J.W.S., S.T.S., G.M.S. **Project administration:** J.W.S., S.T.S., G.M.S. **Funding acquisition:** S.S., J.P.G., P.R.L. S.T.S., G.M.S. **Writing – original draft:** N.S.G., J.J.R., S.S., G.M.S. **Writing – review & editing:** N.S.G., J.J.R., S.S., J.J.A., P.L., R.L.P., Y.R., K.A.T., J.M.S., T.A.A., J.W.C., A.A.M., J.G.S., S.A.S., A.J.D., D.S.R., L.K., M.V., J.P.G., P.R.L., S.B., K.N.C., J.O.W., J.W.S., S.T.S., G.M.S.

### Declaration of generative AI and AI-assisted technologies

During the preparation of this work, the authors used Microsoft Copilot to improve readability and grammar, and for communication support. After using this tool, the authors reviewed, edited, and verified all content and take full responsibility for the content of this publication.

### Data Availability

Human breast cancer data from the METABRIC invasive breast carcinoma cohort were obtained from cBioPortal at <u>cBioPortal for Cancer Genomics</u>. Data generated during this study are available upon request to the corresponding author.

## Notes

### Competing Interest Statement

The authors have declared no competing interest.

