## Supplemental Figures and Legends for "Breast Cancer-derived PDGFB Remodels the CNS Vascular Niche to Promote Brain, Spinal and Leptomeningeal Metastasis"

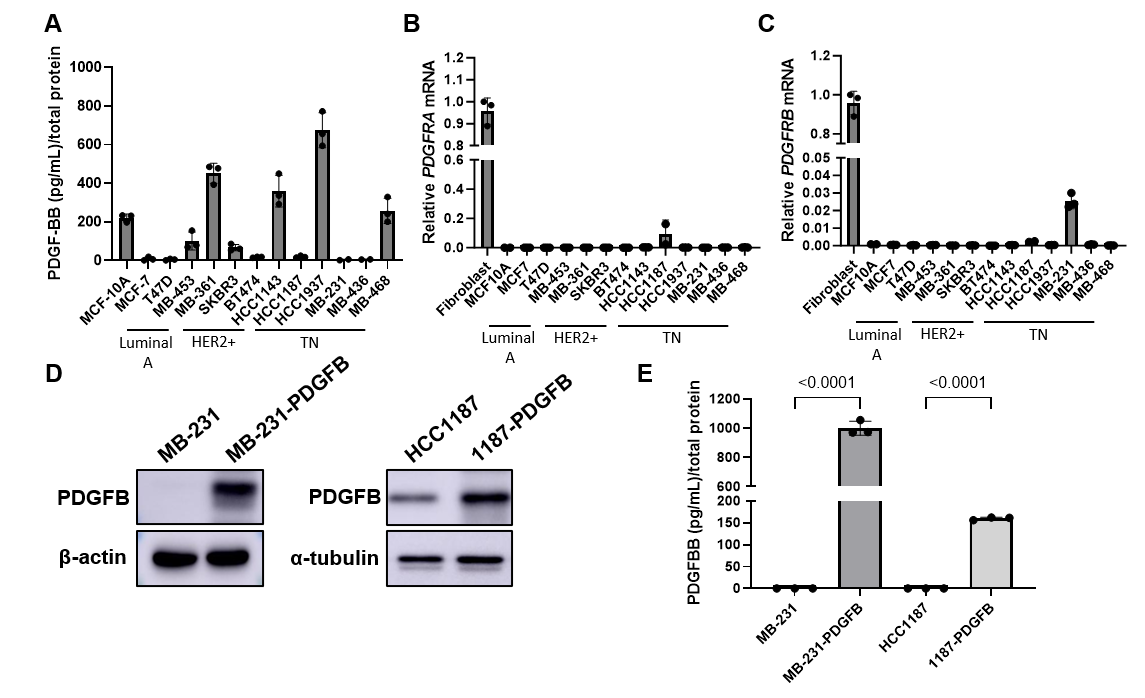
**Supplemental Figure 1. PDGFB signaling components are differentially expressed in BC.**

**(A-B)** Quantitative RT-PCR (qRT-PCR) analysis of (A) *PDGFRA* and (B) *PDGFRB* mRNA expression in a panel of human breast cancer cell lines relative to primary human breast fibroblasts. Gene expression was normalized to *GAPDH*. Data are presented as mean of biological replicates ± SEM (n=3).

**(C)** Quantification of secreted PDGF-BB protein from breast cancer cell lines as measured by ELISA and normalized to total cellular protein content. Data are presented as mean of biological replicates ± SEM (n=3).

**(D)** Western blot validation of PDGFB protein overexpression in MB-231 and HCC1187 cells transduced with either control or PDGFB-expressing lentivirus.

**(E)** Quantification of secreted PDGF-BB from MB-231, MB-231-PDGFB, HCC1187, and 1187-PDGFB cells as measured by ELISA and normalized to total cellular protein content. Data are presented as mean of biological replicates ± SEM (n=3).

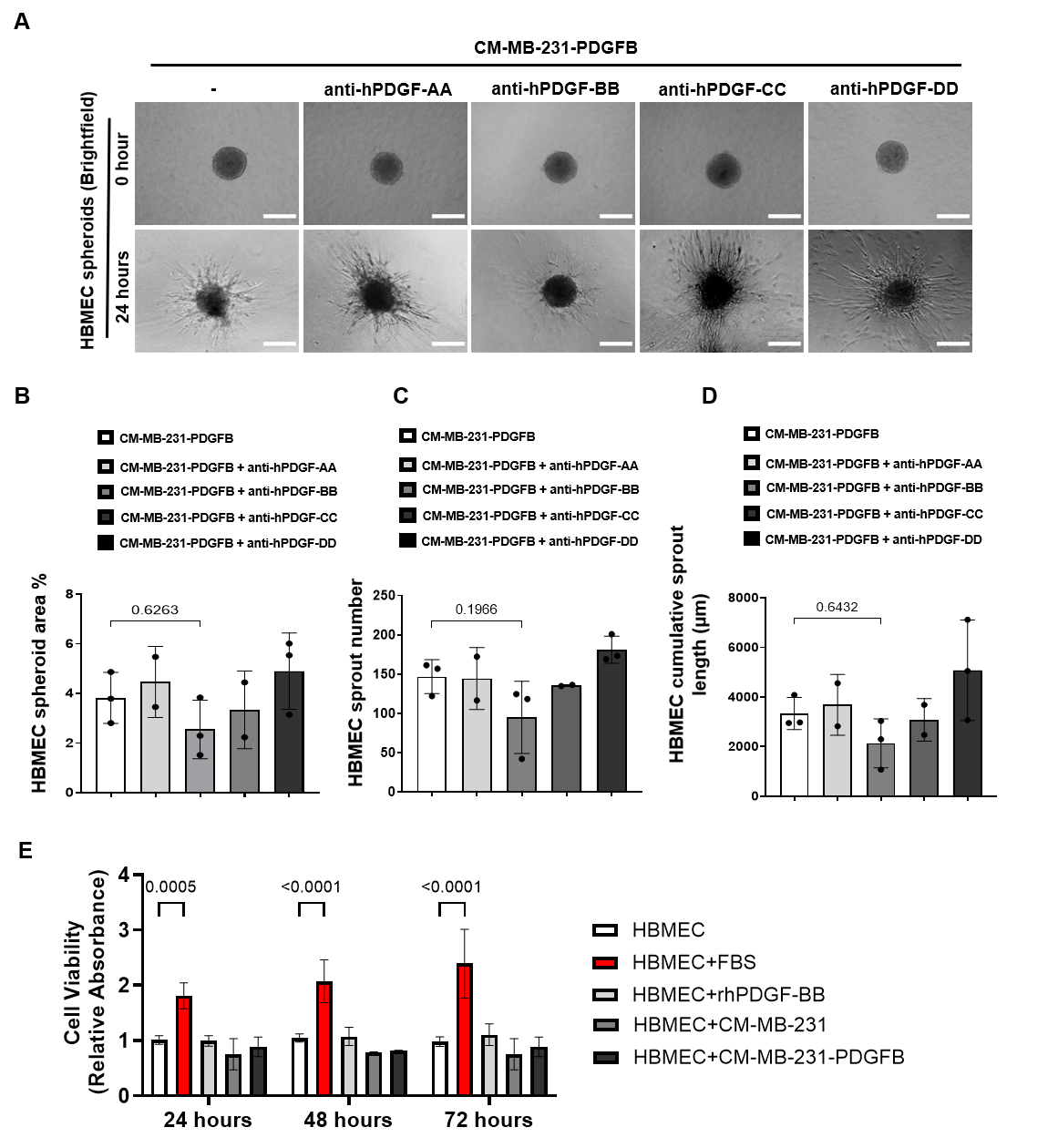

**Supplemental Figure 2. Tumor-derived PDGFB contributes to HBMEC sprouting but not endothelial proliferation.**

**(A)** Representative images of 3D HBMEC spheroids treated with CM-MB-231-PDGFB alone or CM-MB-231-PDGFB supplemented with neutralizing antibodies against PDGF-AA, PDGF-BB, PDGF-CC, or PDGF-DD. Images were acquired after 24 hours. Scale bar, 100 μm.

**(B-D)** Quantification of (B) spheroid area (%), (C) sprout number, and (D) cumulative sprout length. Data are presented as mean ± SEM from biological replicates (n=3).

**(E)** HBMEC viability following treatment with complete growth media (FBS), recombinant human PDGF-BB (100 ng/mL), CM-MB-231, or CM-MB-231-PDGFB for 24, 48, and 72 hours, measured by MTT assay. Data are presented as relative absorbance and shown as mean ± SEM from biological replicates (n=3).

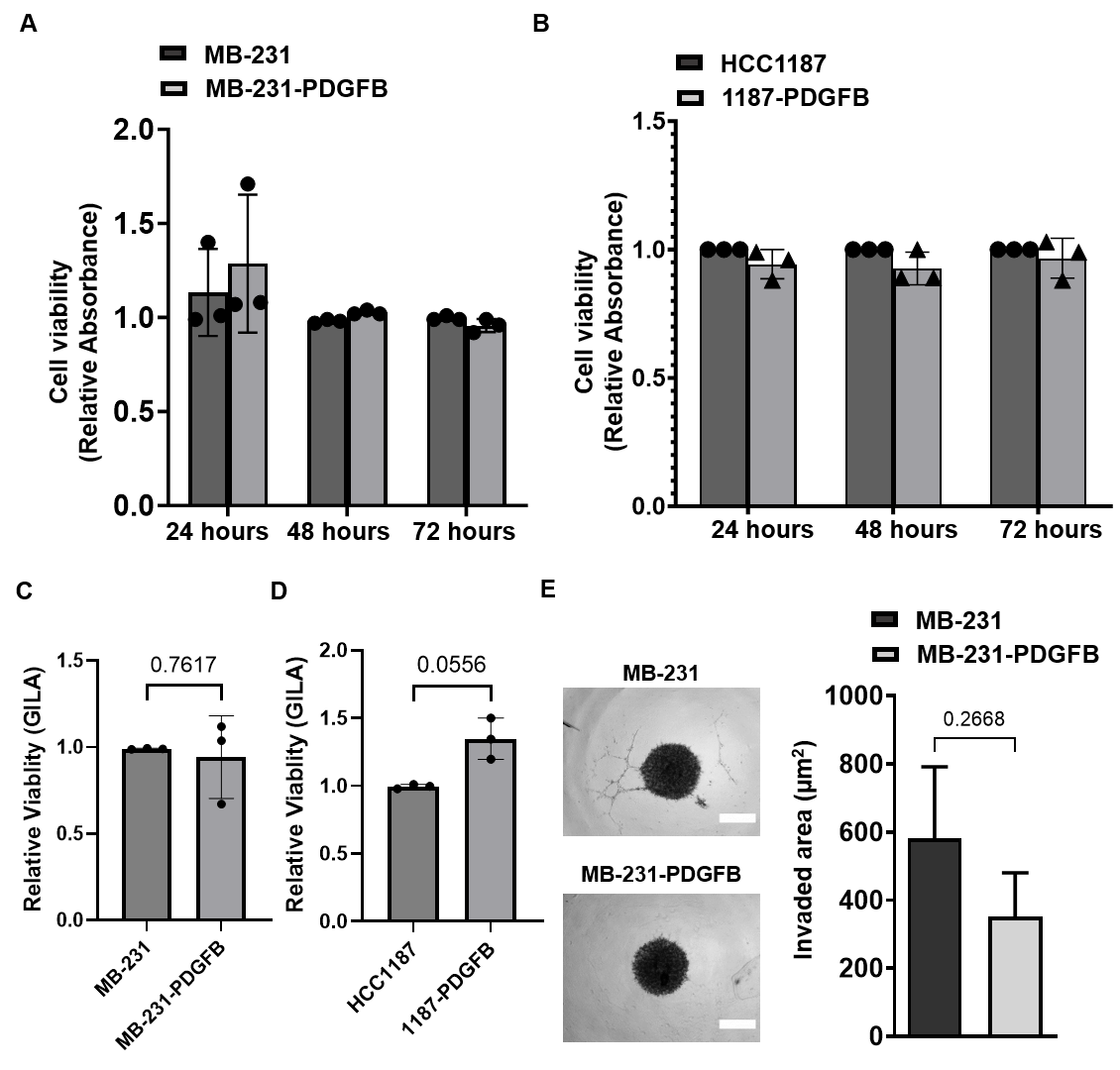
**Supplemental Figure 3. PDGFB overexpression does not alter breast cancer cell viability or invasion *in vitro*.**

**(A-B)** Effect of PDGFB overexpression on (A) MB-231 and (B) HCC1187 cell viability measured by MTT assay at 24, 48, and 72 hours. Data are presented as relative absorbance normalized to the 24-hour control condition and shown as mean ± SEM from biological replicates (n=3).

**(C-D)** Effect of PDGFB overexpression on (A) MB-231 and (B) HCC1187 cell viability measured by GILA assay. Data are presented as relative luminescence normalized to control and shown as mean ± SEM from biological replicates (n=3).

**(E)** Representative images and quantification of 3D spheroid invasion by MB-231 and MB-231-PDGFB cells embedded in basement membrane extract and monitored over 4 days. Invaded area was quantified by ImageJ. Data are presented as mean ± SEM from biological replicates (n≥3 spheroids per condition). Scale bar, 20 μm.

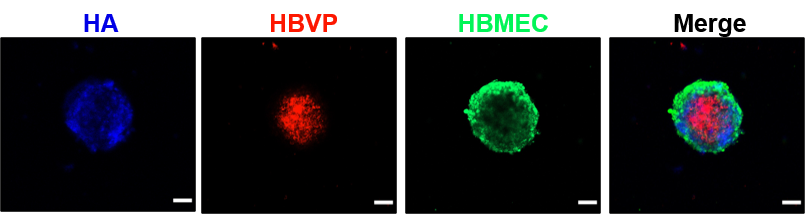
**Supplemental Figure 4. Assembly and organization of tri-culture BBB spheroids.**

Representative confocal images of BBB spheroids composed of human astrocytes (HAs; blue), human brain vascular pericytes (HBVPs; red), and human brain microvascular endothelial cells (HBMECs; green). Images demonstrate organization of astrocytes within the spheroid core, surrounded by pericytes and an outer endothelial layer (i.e. outside of spheroid indicates within brain vessel). Z-stack confocal imaging was performed to confirm cellular distribution throughout the spheroid structure. Scale bar, 50 μm.

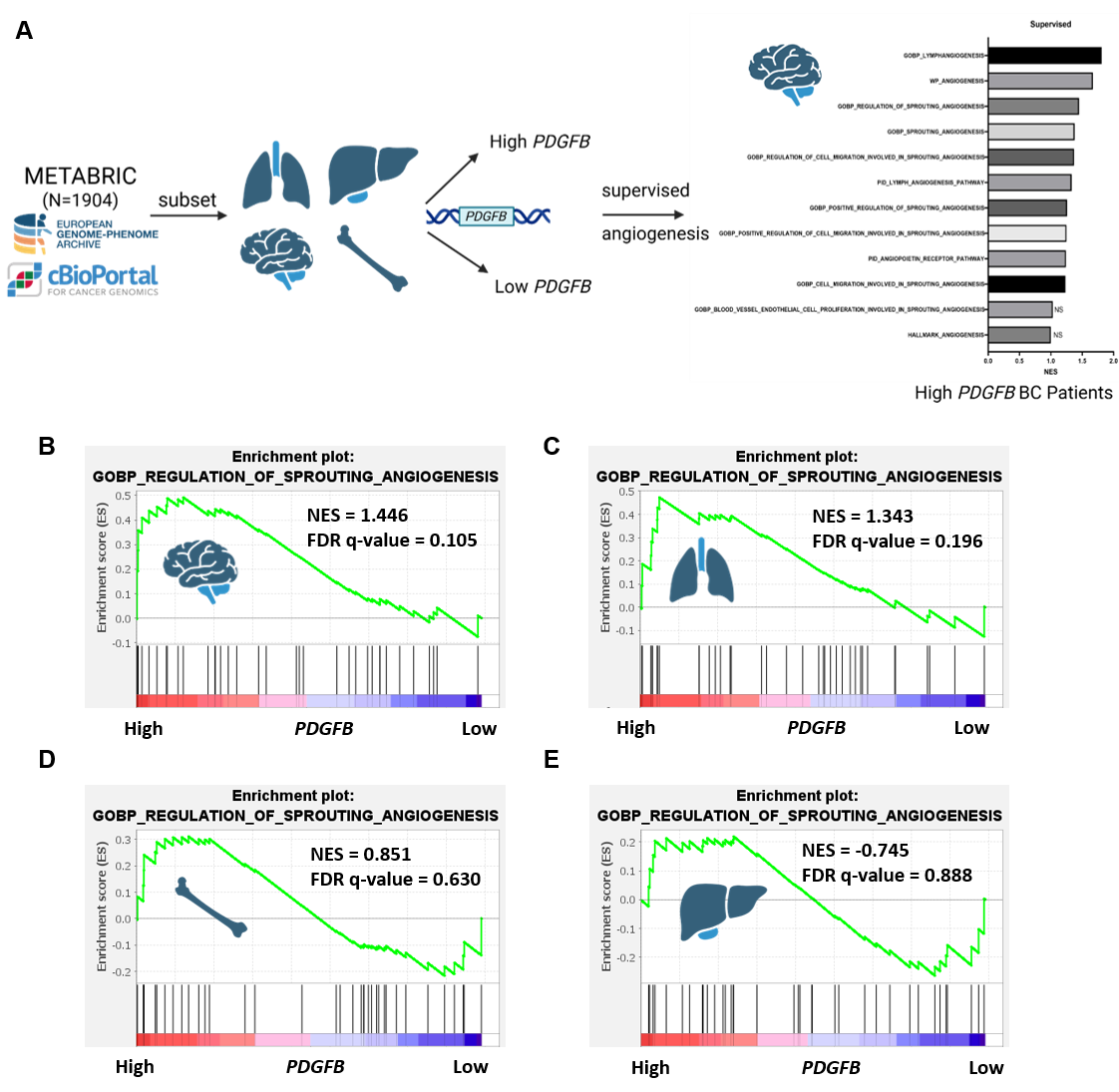
**Supplemental Figure 5. Supervised gene set enrichment analysis results.**

**(A)** Schematic of METABRIC dataset analysis workflow. Primary breast tumors from patients who subsequently developed metastasis were stratified into *PDGFB*-high and *PDGFB*-low groups and analyzed by Gene Set Enrichment Analysis (GSEA).

**(B–E)** Representative GSEA plots of angiogenesis-associated gene sets in patients that developed (B) brain, (C) lung, (D) bone, and (E) liver metastases.
