## Supplementary material for "Breast Cancer-derived PDGFB Remodels the CNS Vascular Niche to Promote Brain, Spinal and Leptomeningeal Metastasis": Supp Methods

**Cell lines**

Cell lines with culture conditions are listed in **Suppl. Table 1.** Primary human breast fibroblasts were cultured as described previously^1^. The mouse TNBC cell line DB7 *Pdgfb* knockdown/control cells were cultured and transduced as previously described^2^. Both human breast cancer cell lines MDA-MB-231 and HCC1187 were transduced using lentivirus containing firefly luciferase obtained from VectorBuilder (#VB900088-2567cxn) and subsequently subjected to selection using geneticin (Fisher, #11811023) for at least 10 days before use. The luciferase transduced cells were infected with lentivirus carrying the human *PDGFB* open reading frame (ORF) from OriGene Technologies Inc. (#RC206587L4V), or a vector control from OriGene (#TR30021V). Cells transduced with control and *PDGFB*-ORF lentivirus were subjected to puromycin selection (Sigma, #P8833) for more than 10 days before use. Primary cell lines were utilized during cell passages two to five.

**Preparation of conditioned medium**

Cells were seeded in 10 cm^2^ dishes at a concentration of 2 x 10^6^ cells and grown in medium containing 10% FBS. Following 48 hours, the medium was removed, and the cells were rinsed with phosphate buffer saline (PBS; pH 7.4). Five milliliters of endothelial basal media with a concentration of 0.1% FBS was introduced, and the cells were incubated for 24 hours at 37°C with 5% CO_2_. The conditioned media (CM) obtained from cells was collected and subjected to centrifugation at 2000 RPM for 10 minutes at 4°C to eliminate cell debris. The resulting supernatant was collected and immediately utilized or stored at -80°C until needed.

**Quantitative real-time PCR**

TRIzol reagent (ThermoFisher Scientific, #15596018) was used to extract total RNA following the guidelines provided by the manufacturer. DNase treatment and cDNA synthesis was conducted using the Maxima First Strand cDNA Synthesis Kit for qRT-PCR (ThermoFisher Scientific, #K1672). The Applied Biosystems StepOnePlus real-time PCR machine and software were utilized to evaluate relative gene expression via human *PDGFRB* (Hs01019589_m1), *PDGFRA* (Hs00998018_m1), and *GAPDH* (Hs99999905_m1) TaqMan gene expression assays.

**Western blotting and immunodetection**

Protein was collected as described previously^3^. Protein lysates were quantified using the Pierce™ BCA Protein Assay Kit (ThermoFisher, #23225), separated using sodium dodecyl sulfate-polyacrylamide gel electrophoresis (SDS-PAGE), and transferred to a polyvinylidene difluoride membrane (PVDF, Biorad, #1620177). PVDF membranes were blocked in 1X TBST containing 5% bovine serum albumin (BSA, Research Products International, #A30075). Membranes were incubated overnight with primary antibodies in 1X TBST with 5% BSA overnight and secondary antibodies for 1 hour. Antibodies are listed in **Suppl. Table 2**. Signal was detected via a GE Amersham Imager 600 (GE Healthcare Biosciences) using ECL detection reagents (Millipore-Sigma, #WBKLS0500).

**3D spheroid invasion assay of breast cancer cells**

MB-231 and MB-231-PDGFB spheroids were generated using a protocol from Andersen and colleagues with minor adjustment^4^. Briefly, cells were collected and placed in a mixture of DMEM and 1X Spheroid Formation Matrix (Trevigen, #3500-096-SP). Cells were seeded at 5 x 10^3^ cells per 50 µL in a round bottom ultra-low attachment (ULA) 96 well plate (Corning, #CLS7007-24EA). For spheroid formation, ULA plates were subjected to centrifugation at 750g for 10 minutes at 4°C. Subsequently, they were incubated at 37°C with 5% CO_2_ for a period of 7 to 10 days to allow for spheroid formation. For this experiment, the spheroids were moved to a fresh ULA 96 well plate and placed into Cultrex Basement Membrane Extract (BME) (Trevigen; #3432-001-01). The spheroids were placed at the center via centrifugation at 300 g for 5 minutes at 4°C. They were then incubated for 1 hour at 37°C with 5% CO_2_. 100 µL of pre-warmed RPMI media was added gradually in drops to the well holding the spheroids and incubated for 4 days. The spheroids were observed using the EVOS Core XL microscope at a magnification of 4X regularly to track cell invasion. The degree of invasion was quantified by determining surface area occupied by the spheroids, with a minimum of three spheroids per experimental group, using ImageJ software.

**3D endothelial spheroid and angiogenesis assays**

3D HBMEC and HUVEC spheroids were generated using the method previously reported by Shah and colleagues, with slight modifications^5^. HBMECs and HUVECs were detached using trypsin and then suspended in a basal medium for endothelial cells (ECM), which included 20% methylcellulose (Sigma, #M0430) and 5% FBS at 1 x 10^3^ cells per 25 μL. Using a hanging drop approach, 25 μL of cell suspension was deposited onto the lid of a 15 cm^2^ culture plate, which was then flipped over and placed onto the bottom chamber which was filled with PBS. The droplets were cultured for 24 hours at 37°C and 5% CO_2_ to create spheroids of consistent size. Subsequently, spheroids were gathered and reconstituted in basal ECM solution containing 5% FBS along with 40% methylcellulose to hinder settling. The spheroid suspension was mixed with neutralized type I collagen (Corning, #354236) and immediately placed on pre-warmed 24-well plates. The mixture was then allowed to solidify at 37°C with 5% CO_2_. After 30 minutes, 1 mL of basal ECM, with or without recombinant human platelet-derived growth factor B (rhPDGF-BB, 100 ng/mL, R&D #220-BB-010), with or without recombinant human vascular endothelial growth factor A (rhVEGFA, 50 ng/mL, R&D #293-VE), or 1 mL of CM, was introduced onto the gel. Subsequently, the plate was placed in an incubator for 24 hours. Spheroids were then observed under an EVOS Core XL microscope at a magnification of 20X to monitor the formation of sprouts. For analysis, five images from each group were batch processed in RStudio (R version 4.1.0) using the ‘SPE’ function in the ‘imager’ package to generate a homogeneous background and further thresholded using the ‘thresh’ function to create binary, black and white images as previously described^6^. The sphere region was outlined and subtracted from the thresholded image and the remaining cell clusters or sprouts were analyzed for total number of sprout as well as length based on Feret diameter using the ‘Analyze Particles’ function in ImageJ^7^. The cumulative sprout length was measured by adding the length of each sprout.

**Development of multicellular co-culture BBB spheroids**

The formation of multicellular co-culture BBB spheroids was carried out according to the protocol described by Choi-Fong Cho *et al*.^8^. In brief, a low adhesion 96-well plate was created by applying a layer of 1% (w/v) agarose (ThermoFisher Scientific, #17852) to the surface of the plate. Three immortalized human BBB cell lines (HBMECs, HAs, and HBVPs) were plated sequentially using BBB working media (basal ECM with 2% FBS) with each cell line being seeded at an equal ratio of 1:1:1 with 3 x 10^3^ total cells. The cells were incubated for 48 hours to facilitate the formation of BBB spheroids.

**Immuno-fluorescent imaging of live cells in BBB spheroids**

In order to examine the distribution of different cell types within the BBB spheroids, HBMECs, HBVPs, and HAs were pre-labeled with PKH67 green fluorescent cell linker kit (Sigma, #MIMI67), PKH26 red fluorescent cell linker kit (Sigma, #MINI26), or CellTracker dye (ThermoFisher Scientific, #C2110) respectively. The BBB spheroid, which contained these pre-labeled cells, was carefully collected and rinsed with PBS three times. Afterward, the BBB spheroids were treated with 3.7% PFA for 10 minutes at room temperature and then moved to a NuncTM Lab-TekTM II chambered cover glass (ThermoFisher Scientific, #12-565-338) for imaging. Confocal Z-stack pictures were obtained for each spheroid, with each slice having a thickness of 10 µm. The images covered a depth of up to 120 µm. The 20X photos were captured using a Zeiss LSM800 confocal microscope.

**Cell migration assay in BBB spheroids**

Approximately twenty BBB spheroids were placed in each well of 6 well culture plates. The spheroids were treated with either vehicle control, rhPDGF-BB (R&D, #220-BB-010) at a concentration of 100 ng/mL, or CM derived from breast cancer cells for 24 hours. The spheroids were incubated at a temperature of 37°C with 5% CO_2_ for 24 hours and then were imaged using an EVOS Core XL microscope with a 10X objective lens to track cell movement. For the analysis, five pictures were processed using the HBMEC and HUVEC 3D spheroid formation and angiogenesis assay method described above. Cell migration was assessed based on their migration capacity or area fraction, which refers to the total area covered by the cells migrating outwards from the sphere relative to the area of the visible frame. To measure the area of only the migrating cells, the sphere region was outlined and subtracted from the thresholded image and the remaining cell clusters were analyzed for area of each cluster and area fraction using the ‘Analyze Particles’ function in ImageJ as previously described^6^.

**Dextran permeability assay in BBB spheroids**

HBMECs, HAs, and HBVPs were grown together for 48 hours, following the previously described method to create multicellular 3D BBB spheroids. Twenty spheroids were cultured with either vector control, rhPDGF-BB (100 ng/mL), or breast cancer-derived CM for 24 hours. Additionally, 50 µg/mL of Tetramethylrhodamine isothiocyanate–Dextran 155 kDa (Sigma, #T1287) was added to the basal endothelial medium. The spheroids were rinsed with PBS three times and then moved to NuncTM Lab-TekTM II chambered cover glass. Each spheroid was imaged with a 20X magnification, capturing confocal Z-stack pictures at 10 µm intervals up to a depth of 120 µm. The mean fluorescence intensity was quantified at the central region of the spheroid, namely at a depth of 70 µM, using Image J. Mean intensity was normalized to spheroid area.

**Cell viability assay**

A MTT kit from Millipore-Sigma (#11465007001), was employed to measure cellular viability. In summary, a total of 6 x 10^3^ cells were placed in a 96 well-plate and given 24 hours to adhere in a suitable culture medium. On the subsequent day, the cells were rinsed with PBS and treated with breast cancer-derived CM for durations of 24, 48, and 72 hours. A solution of MTT at a concentration of 0.5 mg/mL was introduced into each well and allowed to incubate for 4 hours at a temperature of 37°C. A solubilization agent of 100 µL was added and the plates were incubated overnight. The optical density at 550 nm was measured using a Promega GloMax Discover Microplate Reader (GM3000).

**Enzyme-Linked immunosorbent assay**

The levels of secreted PDGF-BB in the supernatant of both control cells and cells overexpressing PDGFB were measured using a human PDGFBB-ELISA kit (R&D, #DBB00) according to the instructions provided by the manufacturer. The resultant PDGF-BB protein concentration was normalized by total lysate protein concentration from the same plate of cells.

**Microfluidic device fabrication and permeability assay**

Microfluidic lumen devices were prepared as previously described^9^. After fabrication, microdevices were pre-coated with 2mg/mL poly(dopamine) to enhance collagen gel attachment to poly(dimethylsiloxane)^10,11^. Type I collagen and hyaluronic acid were added to the central ECM chamber at concentrations of 3 mg/mL (Corning, #354249) and 1mg/mL (Sigma-Aldrich, #H7630), respectively. This hydrogel was casted around a 250 µm diameter nitinol wire that ran from inlet to outlet port. Following gel polymerization, the wire was removed, leaving a hollow cylindrical lumen encased within the hydrogel. After casting, lumens were coated overnight at 37°C with 100uL of 100µg/mL fibronectin (Fisher Scientific, #FC010) in Attachment Factor (Cell Systems, #AZ0-201). HBMECs (Cell Systems) were cultured to 85% confluency before harvesting using Passage Reagent Group (Cell Systems, #4Z0-800). Passages 6-8 were used for all experiments. The following morning, 1 x 10^5^ HBMECs were added to each empty lumen. Devices were then placed in humidified dishes and flip steps were completed for 1.5 hours (30 mins upside-down, 15 min on each side, 20 minutes right-side-up, and 10mins upside-down). Non-adhered cells within lumens were removed and fresh media was added. Additionally, gel-loading lateral ports were covered with bio-inert tape (3M) to minimize evaporation effects and improve barrier integrity. Devices were then cultured overnight within humified dishes at 37°C with 5% CO_2_. From Day 1- 2, lumen devices were imaged and cultured with normal Lonza EGM-2 media or CM from MB-231-PDGFB/1187-PDGFB and respective controls. Fluorescent tracer (20 kDa FITC-Dextran, Invitrogen #D1820) was resuspended in cell culture media at a concentration of 12.5mg/mL. On Day 2, after clearing and imaging the vessel, 50mL of FITC-dextran solution was introduced into the empty inlet port of the lumen device. Images were taken every 5 seconds for 120 seconds using a 5X objective on a Nikon AX confocal microscope to track vessel permeability. Time-lapse images were then recorded and processed with a custom MATLAB code used to calculate apparent vessel permeability, defined as the net rate of tracer efflux across the endothelial wall into the surrounding ECM^12^.

**Immunostaining of HBMECs**

HBMECs were plated at a density of 1×10^5^ cells on glass coverslips and maintained in complete culture medium. After reaching a confluency of 90-100%, they were exposed to breast cancer-derived CM for 24 hours, rinsed and fixed with ice-cold methanol for 15 minutes. Fixed HBMECs were then blocked in 1X PBS, 5% normal donkey serum, and 0.3% TritonTM X-100 for 1 hour. Subsequently, cells underwent three washes and were then subjected to overnight primary antibody incubation at 4°C. On day 2, the cells were rinsed three times with 1X PBS and then incubated with secondary antibody for 1 hour, while being shielded from light. All antibodies are listed in **Suppl. Table 2**. The cells were stained with Hoechst dye to visualize nuclei. The samples were mounted and imaged using the EVOS M-7000 microscope.

**Xenograft and Allograft Experimental Metastasis Studies**

Female nod scid gamma (NSG) were obtained from the OSUCCC Preclinical Therapeutics Mouse Modeling Shared Resource; breeders for the colony were received from Jackson Labs strain #005557 (RRID:IMSR_JAX:005557). Female FVB/N mice were obtained internally through established breeders (Jackson Labs, #001800, RRID:IMSR_JAX:001800). Upon the first mouse reaching early removal criteria (ERC) for each injection model, animals were euthanized via CO_2_ inhalation and perfused with PBS followed by 4% PFA. Brains were fixed overnight in 4% PFA and then swapped to 30% sucrose for an additional 48 hours. Brains were cryoembedded in OCT and 15µm sections were obtained on Leica cryostat. The remaining organs were fixed in formalin. Spines were further decalcified and paraffin embedded at the OSUCCC Comparative Pathology and Digital Imaging Shared Resource.

**Bioluminescent IVIS imaging**

Bioluminescent imaging was performed using the PerkinElmer IVIS Spectrum 2 imager in the OSUCCC Small Animal Imaging Shared Resource. The mice were anesthetized using 2% isoflurane (OSUWMC Main Pharmacy, #NDC66794-017-10) and then intraperitoneally injected with 200 µL of 150 mg/kg D-luciferin (Goldbio, #eLUCK-100). After 8 minutes, images were taken using both an auto-exposure and a 60-second exposure time until they reached max intensity. The Living Image program (Revvity) was used to quantify the intensity of bioluminescent signals.

**Evans Blue Permeability Studies**

For the HCC1187 cohort, upon the first mouse of each cohort reaching ERC, 2% Evans blue solution was administered I.P. at 4 mL/kg, and allowed to circulate for 2 hours. Mice were euthanized via CO_2_ inhalation and perfused with PBS followed by 4% PFA. Brains were fixed for 2 additional hours in the absence of light and cryoprotected for 48 hours in 30% sucrose. Brains were cryoembedded and sections were evaluated for intratumoral Evans blue uptake by 647nm fluorescence. For the DB7 cohort, a 2% Evans blue solution was administered I.P. at 4 mL/kg and allowed to circulate for 2 hours. Mice were euthanized via CO_2_ inhalation and perfused with PBS. A 2.5 cm spinal section was taken for histological evaluation from each mouse, fixed for 2 hours in 4% PFA, decalcified in 0.5M EDTA for 48 hours, and cryoprotected for 48 hours in 30% sucrose and frozen in OCT. The remaining spinal cord was weighed and then immediately placed into *N,N*-Dimethylformamide for 48 hours at 55°C. Extracted Evans blue was then quantified via absorbance at 600nm. Brains were processed as described above. 15 µm sections were obtained and analyzed at 647nm for Evans blue uptake.

**Immunostaining of Frozen Sections**

15 µm cryosections were stored at -80°C until ready for staining. Slides were fixed with 4% PFA for 15 minutes, washed with 1X PBS, and blocked in a solution of PBS with 10% NDS, 5% BSA, and 0.3% TritonX100 (MilliPore Sigma, X100) for 1 hour at RT. Slides were treated with Mouse on Mouse (Vector Laboratories MKB-2213-1) block for an additional hour when appropriate. Subsequently, slides were subjected to overnight primary antibody incubation at 4°C. The slides were rinsed three times then incubated with secondary antibody for 2 hours, while being shielded from light. All antibodies are listed in **Suppl. Table 2**. The sections were stained with Hoechst dye to visualize nuclei. The samples were mounted with ProLong Gold antifade (Invitrogen #P10144) and imaged using the EVOS M-7000 microscope. Immunostaining was quantified as previously described^13^.

**Immunostaining of Paraffin sections**

Sections were baked overnight at 65°C and washed with xylenes for deparaffinization. EDTA antigen retrieval (pH 9.0) was performed at 110°C for 10 minutes. Samples were blocked as described. Subsequently, slides were subjected to overnight primary antibody incubation at 4°C. On day 2, the slides were rinsed three times and then incubated with secondary antibody for 2 hours, while being shielded from light. All antibodies are listed in **Suppl. Table 2**. The sections were stained with Hoechst dye to visualize nuclei. The samples were mounted with ProLong Gold antifade (Invitrogen #P10144) and imaged using the EVOS M-7000 microscope. Immunostaining was quantified as previously described^13^.

**Histopathologic analysis of spinal metastasis**

Fixed tissues were routinely processed on a Leica Histocore Peloris Tissue Processor (Leica Microsystems), embedded in paraffin wax, sectioned by a certified histotechnologist, stained with Hematoxylin & Eosin (H&E) and coverslipped on a Leica HistoCore SPECTRA and CV Autostainer/Automated Coverslipper (Leica Microsystems). A board-certified comparative veterinary pathologist (Corps, K.) with greater than 12 years of experience examined two separate cohorts of slides. Photomicrographs were taken on a Nikon Eclipse Ci-L light microscope (Nikon Instruments Inc.) with an attached Olympus camera and cellSens Software (Olympus/Evident Scientific).

**Analysis of human breast cancer patient data**

Human breast cancer patient data from the METABRIC invasive breast carcinoma cohort (N = 1904) was obtained from cBioPortal. Patients with lung (N = 131), liver (N = 152), brain (N = 66) and bone metastases (N = 259) were included in downstream analysis. Analysis was conducted in R 4.2.2 utilizing dplyr 1.0.10, tibble 3.1.8, ggplot2 3.4.0, and ggfortify 0.4.15 packages to import, subset, and analyze relevant patient data. Gene expression data for all patients with lung, liver, brain, or bone metastases were each divided into two groups by median *PDGFB* expression and exported for use in Gene Set Enrichment Analysis (GSEA). Unsupervised GSEA was conducted using GSEA 4.2.3 for windows for all available datasets from the C2 (Curated Gene Sets) collection, which identified WP_ANGIOGENSIS as a significantly enriched gene set in high *PDGFB* brain metastasis patients. Supervised GSEA was then performed for select angiogenesis-related gene sets (Supplemental Table 1) for all patients from the human Molecular Signatures Database. Gene sets with nominal p-value < 0.05 (unsupervised analysis) and FDR q-value < 0.25 (supervised analysis) were considered significantly enriched per the Broad Institute guidelines.

**Statistical analysis**

GraphPad Prism 10.6.0 was used for conducting statistical analysis. Sample sizes were not predetermined statistically. Prior to analysis, data were checked for normality via Shapiro–Wilk, Kolmogorov–Smirnov and D'Agostino–Pearson omnibus normality testing. Data were considered normally distributed after passing at least one of the three tests. The appropriate homoscedastic or heteroscedastic unpaired two-tailed or one-tailed Student t-test was used to compare two groups of normally distributed data. The Mann-Whitney U test was used for statistical comparisons of data that were not normally distributed. For analyses involving three or more groups, a one-way ANOVA was used and pair-wise comparisons were computed post-hoc with appropriate correction. Statistical significance was established at *P* ≤ 0.05.

**Supplemental Methods References**
